# Transcriptome analyses of tumor-adjacent somatic tissues reveal genes co-expressed with transposable elements

**DOI:** 10.1101/385062

**Authors:** Nicky Chung, G.M. Jonaid, Sophia Quinton, Austin Ross, Corinne E. Sexton, Adrian Alberto, Cody Clymer, Daphnie Churchill, Omar Navarro Leija, Mira V. Han

## Abstract

**Background:** Despite the long-held assumption that transposons are normally only expressed in the germ-line, recent evidence shows that transcripts of transposable element (TE) sequences are frequently found in the somatic cells. However, the extent of variation in TE transcript levels across different tissues and different individuals are unknown, and the co-expression between TEs and host gene mRNAs have not been examined.

**Results:** Here we report the variation in TE derived transcript levels across tissues and between individuals observed in the non-tumorous tissues collected for The Cancer Genome Atlas. We found core TE co-expression modules consisting mainly of transposons, showing correlated expression across broad classes of TEs. Despite this co-expression within tissues, there are individual TE loci that exhibit tissue-specific expression patterns, when compared across tissues. The core TE modules were negatively correlated with other gene modules that consisted of immune response genes in interferon signaling. KRAB Zinc Finger Proteins (KZFPs) were over-represented gene members of the TE modules, showing positive correlation across multiple tissues. But we did not find overlap between TE-KZFP pairs that are co-expressed and TE-KZFP pairs that are bound in published ChIP-seq studies.

**Conclusions:** We find unexpected variation in TE derived transcripts, within and across non-tumorous tissues. We describe a broad view of the RNA state for non-tumorous tissues exhibiting higher level of TE transcripts. Tissues with higher level of TE transcripts have a broad range of TEs co-expressed, with high expression of a large number of KZFPs, and lower RNA levels of immune genes.

## Background

Although transposable elements (TEs) have been studied for a long time, their ubiquitous and highly tissue-specific expression patterns are starting to be appreciated only recently. The fact that TEs compose close to 40% of the human genome is frequently emphasized, but the fact that there is observable amount of TE derived transcripts in human RNA-seq data has mostly been ignored or regarded as a nuisance without any functional relevance [1]. LINE elements have long been thought to be expressed only in the germline cells [2–4]. But, both full-length and partial transcripts of LINEs are frequently found in the somatic cells [4–6] with large variation in expression levels across tissue types, and among different individuals [7]. The level of TE expression is especially pronounced in cancer cells [8–11], and cell lines [12], but are also observed in neurogenesis [13] and normal somatic tissue. Faulkner et al. in 2009, was the first study to provide a global picture of the significant contribution of retrotransposons to human transcriptome in multiple tissue types [14]. This report showed that 6-30% of transcripts had transcription start sites located within transposons, and these transposons were expressed in a tissue-specific manner and influenced the transcription of nearby genes. The results were extended by Djebali et al. in 2012 showing again the tissue-specificity of transposon expression, and that most of these transcripts are found in the nuclear part of the cell [15]. In addition to the tissue-specific expression of TEs, important regulatory roles for TEs are emerging (reviewed in [16]). Observations include contribution to transcription start sites [14], source of transcription factor binding sites [17], source of long non-coding RNAs [18], active transcription during early development [19], and even critical function similar to long non-coding RNAs that guide chromatin-remodeling complexes to specific loci in the genome [20].

Although there are many reports of TE expression in the somatic cells, there is still a large gap in our understanding of how TE expression is repressed and de-repressed in human somatic cells. Based on what we have learned so far, TE expression is regulated through multiple layers, consisting of transcription factors, epigenetic modification, PIWI-interacting RNAs (piRNAs), RNA interference (RNAi), and posttranscriptional host factors. Recently, two different approaches of genome-wide screening have identified proteins that regulate different aspects of the activities of LINE elements. CRISPR–Cas9 screen was used to identify proteins that restrict LINE activity [21]. The protein MORC2 and the human silencing hub (HUSH) complex was shown to selectively bind evolutionarily young, full-length LINEs located within euchromatic environments, and promote deposition of histone H3 Lys9 trimethylation (H3K9me3) for transcriptional silencing [21]. And through proteomics approaches, two studies have recently identified the localization of ORF1 and ORF2 proteins and its interacting partners [22], and the timing of the entrance of the ORF2 protein complex into the nucleus [23]. But, no study has yet examined the correlation in transcript levels of host mRNA and transposon RNA.

Recently, high-throughput RNA-seq data of various types of cancer samples and their normal counterparts have become available in The Cancer Genome Atlas (TCGA) [24–26]. By focusing on the non-tumorous tissue samples from TCGA, we can access thousands of natural experiments across various types of tissues that show variation in TE transcript levels, and obtain a global picture of TE expression and regulation in humans. An important strength of the TCGA dataset is the large number of samples collected for each tissue type and the high depth of the RNA-seq experiment, with a median of about 150M reads per sample, which is several times larger than a usual RNA-seq library. The variation in TE transcript levels observed in multiple samples within each tissue, allowed us to analyze the co-expression patterns between host genes and TEs for the first time. We hypothesized that genes that regulate the transcription level of TEs would show correlation in expression levels with the TE transcripts. Since the samples are collected from fresh-frozen tissues, TE transcript levels are observed *in vivo*, complementing the studies that focus on retrotransposition assays or transposon expression in human cell lines.

We first summarize the survey of TE expression variation found in the RNA-seq data from 697 samples of cancer-adjacent non-tumorous tissue. We confirm the earlier findings that TE expression varies across tissue types. Transcript levels of individual TE loci are highly tissue specific and within each family only a few individual loci are highly expressed, contributing to the bulk of the transposon transcripts at the family level. We also find large variation in total TE transcript level across individual samples within each tissue type.

Although, transposons have strong tissue-specific patterns at the locus level, we also found that the majority of TEs show global co-expression at the family level across samples. By analyzing the co-expression between these TEs and individual genes, we found co-expression modules of TEs and genes replicated across tissues.

## Results

### TE derived transcripts are quantified across 16 tissues and 697 samples of tumor adjacent controls

We re-aligned and quantified TE derived transcripts from the RNA sequencing data of 697 samples across 16 tissues collected as non-tumorous controls for the TCGA project (supplementary Table 1). The library sizes for these samples range from 50M reads at the minimum, to up to 390M reads, with a median at about 149M reads (75M pairs). Although all tissues included in this study were sequenced using the HiSeq 2000 platform, esophagus and stomach samples were sequenced separately at British Columbia Genome Sciences Centre (BCGSC), with higher sequencing depth on average (median 227M reads). The proportion of reads that do not map to annotated genes were different between the later samples sequenced at University of North Carolina at Chapel Hill (UNC), and the earlier BCGSC sequenced samples, with BCGSC samples having more reads (median 177M) not mapping to annotated genes and discarded, while UNC samples had less reads discarded (median 97M), possibly due to the difference in poly-A enrichment protocol (MultiMACS mRNA isolation kit *vs.* TruSeq RNA Library Prep Kit [27]) among many other differences, including read length. Because of these differences, when comparing across tissue samples, we had to consider esophagus and stomach tissues separately, and they could not be compared against the rest of the tissues.

Despite the differences in the overall sequencing depth and overall proportion of reads mapping to genes, we found that the DESeq2 normalization method normalizes the reads effectively and the correlation due to library size disappears after normalization within tissues (Supplementary Figure 1). Our co-expression analysis is done within each tissue separately. We also replicate our results found in esophagus and stomach with similar results found in at least one other tissue.

Although we find reads mapping to TEs in all the samples that we have examined, the overall transcripts coming from TEs are still a relatively tiny proportion of the total library. The total number of reads mapping to TEs ranged from 137K to 2.1M with a median of 615K for UNC samples, and ranged from 282K to 3.3M with a median of 835K for BCGSC samples. This count excludes some of the potential read-thru transcripts as described in the next section. The total read counts across all TEs amount to about 1.2% (UNC) or 1.7% (BCGSC) of the total reads mapping to known genes in non-tumorous tissue samples.

### TE reads originating from pre-mRNAs or retained introns are corrected by comparing the read depths of the flanking introns

There have been previous reports of transposon reads coming from pre-mRNA or retained introns in the mature RNA of genes that contain TE sequences in their introns [28]. The extent of this problem can be partially estimated by comparing the read depths of the transposon to the read depths of the flanking introns. If the reads mapping to TEs are part of the pre-mRNA or retained introns, we should see continuous mapping of reads that span the introns flanking the TE of interest, and observe reads that map across the intron-TE boundaries. We can also partially correct for this problem by utilizing the read depths in the flanking introns to proportionally reduce the number of total reads mapped to TEs. The approach is described below.

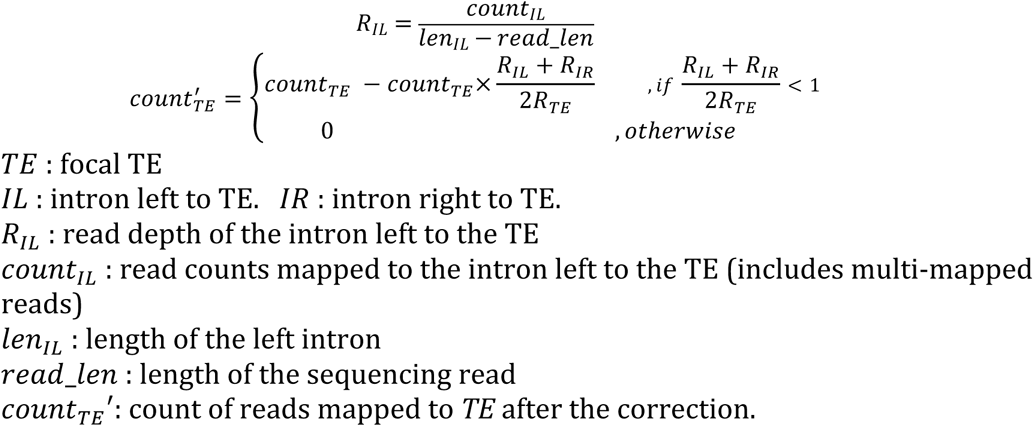

We modified the software TEtranscripts [29], following this approach, to discount the TE read counts based on the read depths of the surrounding introns. By looking for large differences after correcting by flanking read depth, we identified TEs that are most frequently transcribed as part of the introns (Table 1). We also found cases where the method corrected for erroneous TE quantifications due to TEs embedded within long non-coding RNAs (lncRNAs). For example, an AluSx1 element on chromosome Y at position 21153222 (AluSx1_dup59209) had very high transcript levels with an average read count of 18863 in thyroid and head and neck tissue, but the Alu element is embedded in a lncRNA gene called *TTTY14*. The reads mapping here are counted as AluSx1 transcripts based on the UCSC TE annotation, but in the alignment, we see that there are reads spanning the boundaries of AluSx1_dup59209, and almost all the reads mapped in the region are uniquely mapped reads. It looks to be a case of an *Alu* domestication, where an *Alu* insertion or a secondary duplication of an original *Alu* insertion became part of a testis specific RNA gene [30]. Three examples of AluSx1, L2a, and L1MA7, where the read counts for the transposons are reduced to zero are visualized in Supplementary Figure 2. AluSx1_dup59209(chrY:21153222-21153521) is embedded within an exon of gene *TTY14*. L2a_dup21781(chr2:113980079-113981081) is embedded in an intron of *PAX8*. L1MA7_dup4297 (chr8:134015602-134015763) is embedded in an intron of gene *TG*. In all three cases, read counts for the focal TEs were reduced to zero after the correction described above, and the reads mapping to these TEs did not contribute to the overall TE family count. If one is interested in transposable element transcript level that is not part of a longer RNA molecule, it is important to take into account the read depths of the flanking introns or exons, especially the latest non-coding RNA gene annotations, when quantifying repeat element transcripts in the genome.

**Table 1.**
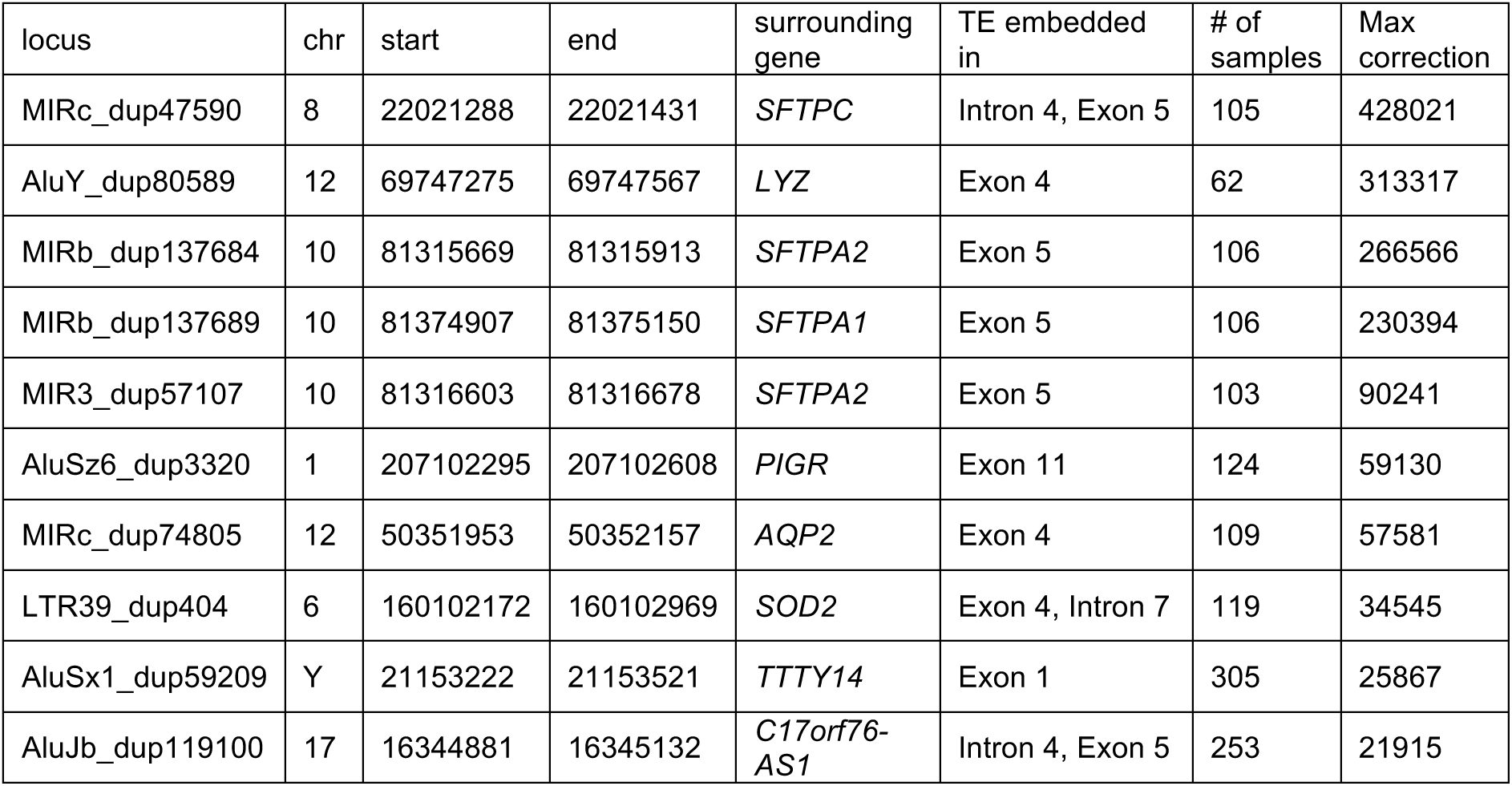
Transposon loci that show large difference after correcting for pre-mRNA/retained introns. a. Transposon loci embedded within introns or exons of genes that frequently result in the largest correction in each sample. Locus id, genomic location, surrounding gene and structure the TE is embedded in, and the maximum number of reads removed in a sample.

### Relying on uniquely mapped reads for repeat quantification results in quantification biased for mappable elements

Due to the difficulty of mapping reads to repeat elements, one of the approaches taken is to count only the reads that map to a unique position in the genome. But this approach has repeatedly been shown to produce results that are worse than expectation-maximization [31, 32], and can lead to serious biases. If we only count uniquely mapped reads in our analysis, not only did we throw away from 10.7% to up to 45% (median 14.2%) of the total TE transcripts, we threw away data in a biased manner, such that we ended up “quantifying mappability” instead of “quantifying transcripts”. This problem is especially pronounced when quantifying the young and active L1HS element. To assess the effect of alignment on quantification, we tried two different alignments, one based on the STAR aligner with up to 200 multi-mapped positions, and the other based on Bowtie1 with only a single best alignment position, discarding all reads that do not have a unique best mapping. Figure 1 shows two cases that illustrate the limitations of either of the approaches. Figure 1 a. shows an example of a full length L1HS locus on chromosome X: 75453754-75459553 with high mappability (48base mappability shown at the top of the panel) due to many accumulated mutations in its sequence. The top panel shows the bowtie1 alignment, allowing only uniquely mapping reads with a single best hit, and the bottom panel shows the STAR alignment with multi-mapping up to 200 mappings per read. In the STAR alignment, we can see erroneously split read alignments at the 3’ end that result in reads mapping across greater than 10K distances, that shows a limitation of a splicing oriented alignment software. The transcription for this element does not start at the 5’ end of the full length, but there is clear and unambiguous transcription starting from about 1500 bases in, that are congruent between both alignments. In Figure 1 b. it shows another full length L1HS locus on chromosome X: 11953208-11959433, this time a young element with very low mappability. Comparing the top and bottom panels, we can see that with the unique mapping we are ignoring all the reads that are perfectly mapping to this locus, but also map to multiple other locations. There is a huge pile-up at the 5’ end of the full length. If we look at the reads mapping to the 5’ end of this locus, their NH tags show numbers ranging from 2 to 4, meaning that they are mapping to two to four alternative locations in the genome. Considering that L1HS loci containing the 5’ ends are more likely to be full length elements, these reads are more likely to be coming from one of the few full length L1HS loci in the genome, but, we end up ignoring these reads if we are only counting uniquely mapping reads. On the other hand, with multi-mapping, we end up quantifying with large uncertainty on whether the reads piled up in this region are really transcribed from this particular locus. This is evident by the small regions of extremely high pile-ups that reflect fragments that are found in the genome with high frequency. Although, we should point out that we don’t count all the reads aligning here at face value, since the expectation maximization algorithm will down weigh the counts of reads by the number of places it maps to.

**Figure 1.**
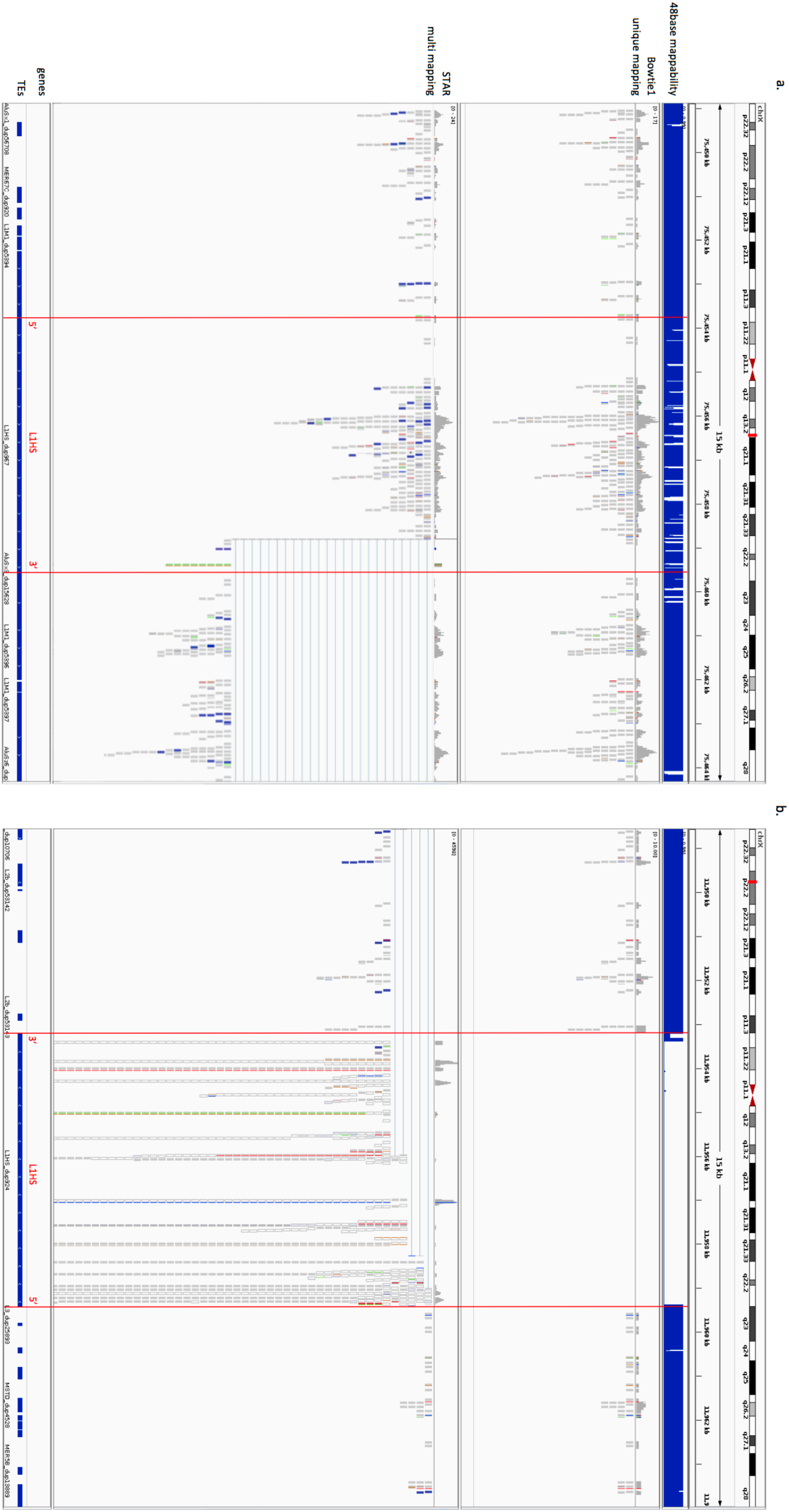
Comparison of read alignment on full length L1HS with multi-mapping and unique mapping. Reads mapped to two different full length L1HS elements from a stomach tissue sample (A4GY) visualized through IGV. a. L1HS_dup967, a 5799nt length element on chromosome X: 75453754-75459553. b. a 6225nt length element L1HS_dup924 on chromosome X: 11953208-11959433. Chromosomal locations, 48 base mappability calculated with GEM, Bowtie1 alignment allowing only uniquely mapped reads with single best hit, STAR alignment allowing multi-mapped reads up to 200 mapping for each read, gene annotation and Repeatmasker TE annotation are shown from top to bottom. Red lines mark the boundary of the L1HS elements with 5’ and 3’ noted.

Comparing the mappability of the two examples, we can see that the uniquely mapping approach preferentially counts reads coming from older TE loci with higher mappability. This can also be shown by correlating the locus level read counts for each L1HS element against the length of uniquely mappable positions in each TE locus (Figure 2 a.). For the unique mapping approach, we see there is significant correlation between the locus level quantification and the total length of uniquely mappable positions within that locus (*p-value* = 5.523e-07 for sum of read counts across all samples, and *p-value* = 1.072e-13 for maximum read count among all samples). The multi mapping approach with Expectation Maximization does not show that bias for uniquely mappable regions (*p-value*= 0.60 for sum and *p-value* = 0.08 for max) (Figure 2 b.).

**Figure 2.**
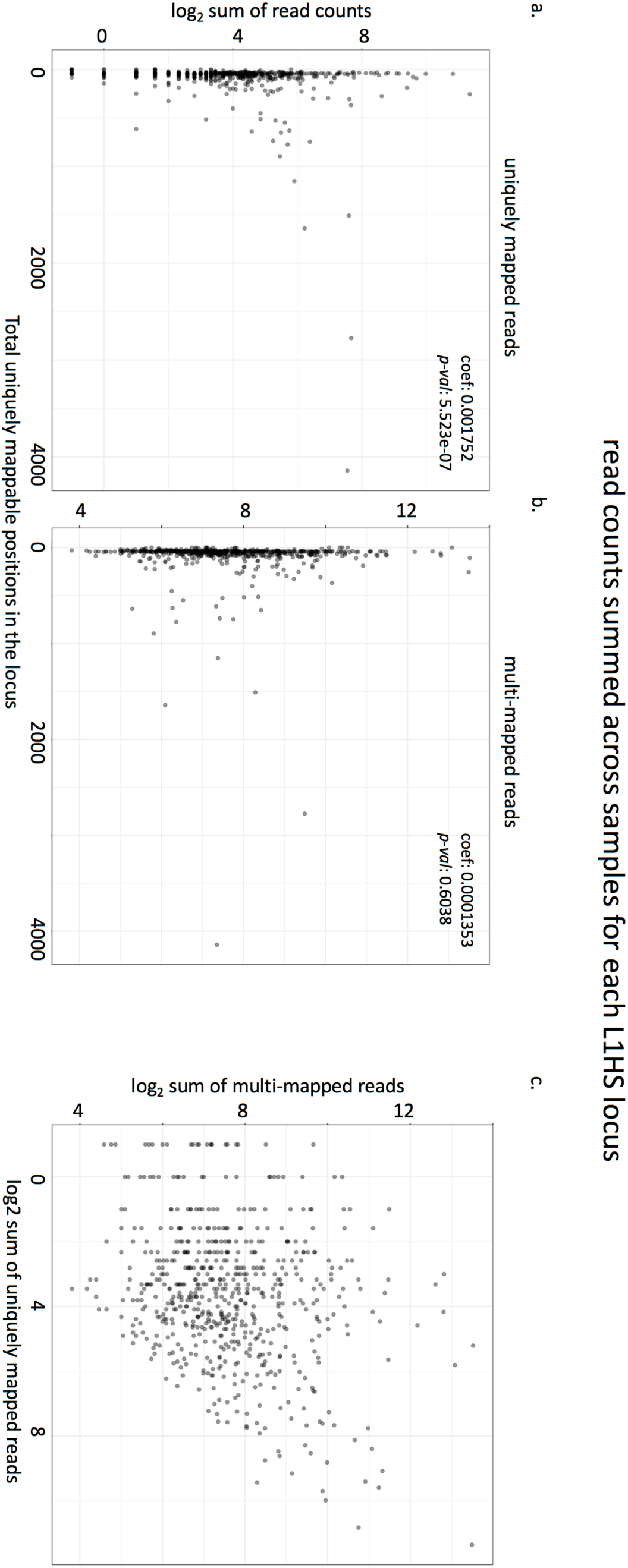
Relationship between read counts for each L1HS locus, and the mappability of the locus. Log_2_ transformed total read counts mapped to each L1HS locus in the hg19 genome, summed across all samples in our dataset are plotted against the total uniquely mappable positions (number of positions with mappability score = 1 based on 48bp mappability calculated with GEM) for each locus. L1HS loci with zero read counts are marked at −1 instead of –infinity. a. read counts from Bowtie1 alignment with uniquely mapped reads only b. read counts from STAR alignment allowing multi-mapped reads up to 200. c. comparison of read counts for each L1HS locus between the uniquely mapped reads and multi-mapped reads.

Consequently, there is limited correlation in the locus level quantification of L1HS between the uniquely mapped reads and the multi-mapped reads (Figure 2 c.). This shows the difficulty of quantifying young active elements, such as L1HS using genome-wide RNA-seq data. Due to these limitations, analysis on L1HS in this study has been done at the family level. The family level quantification of L1HS still shows variability based on the read mapping approach (Supplementary Figure 3 e.), but it shows stronger correlation than the locus level quantification. We still wanted to utilize the abundance of RNA-seq data available for studying L1HS transcription, and glean information on L1HS from these data. Based on the observation that 3’ ends of L1HS are frequently represented in fragmented L1HS loci, while the 5’ ends are more frequent in full length L1HS loci, we decided to use the read counts of the 5’ end of the element as a measure that better represents the transcript level of full-length L1HS transcripts in the sample. All the following analysis on L1HS expression are based on the reads mapped to L1HS sequences in the genome that align with the first 300 bases of the 5’ end of the L1HS consensus sequence and allowing for multi-mapping.

On the other hand, we found that quantification of older elements showed very strong correlation between the two approaches, unique mapping and multi-mapping, reflecting the higher mappability of older elements in the genome (Supplementary Figure 3). For the locus level co-expression analysis with the Zinc Finger Proteins, we limited our analysis to older elements that are 100% uniquely mappable across its sequences with a 48-base read length. All of our main results are qualitatively replicated in the data with uniquely mapped reads aligned with bowtie, except for the results regarding the 5’ end of L1HS expression.

### TE expression shows tissue-specific expression patterns at the locus level among somatic tissues

There have been multiple reports of tissue specific expression of TEs in the human genome, starting from Faulkner et al. in 2009 [14] to Philippe et al. 2017 [12] more recently. We also found highly distinct tissue specificity in TE transcripts in the TCGA data, such that that we could cluster each sample into their broader tissue groupings, based on locus level TE expression patterns alone without relying on any genes at all. Figure 3 shows the clustering of tissues for family level and locus level quantification of LTRs, DNA transposons, SINEs and LINEs. We used normalized mutual information between the different clustering results and the ground truth (the true tissue group) to evaluate the quality of clustering. Normalized mutual information was compared for clustering results based on gene expression, family level TE expression, locus level TE expression and random assignments. We found that the locus-level TE expression was as predictive of tissue groupings as the genes (Figure 3.i, Supplementary Table 2). LTRs, DNA transposons, LINEs and SINEs gave similar clustering accuracy as the genes. The TE family expression levels did not have enough information to cluster the tissues correctly. We note here, that these are loci selected by the rank of variance in log_2_ normalized read count across all samples regardless of tissue type, and we haven’t done any differential expression analysis to identify the markers that are the most informative for accurate tissue classification. Thus, the classification performance we observe here is not the optimal performance that we could get if we were to decide on the markers based on a trained classifier. When we excluded TEs that are within 1K, 10K, and 100K of the start and ends of the genes, the accuracy declined, so part of the tissue specificity is due to co-location with tissue specific genes. But, even when relying on TEs 100K away from any known genes, we saw that tissue specific information was largely retained. On the other hand, when we focused on younger elements, HERVs within LTRs and young L1s within LINEs, there was a large reduction in information content, especially for young L1s. Clustering based on locus level expression for L1HS, L1PA2 and L1PA3 was not any better than clustering based on family level expression of all LINEs. We suspect this is due to the lower locus level mappability and large uncertainty in locus level expression quantification for young L1 elements. Figure 4 shows fifteen representative TE loci that show tissue specific expression. These loci are chosen from TEs that are 100K away from the start or end of any gene.

**Figure 3.**
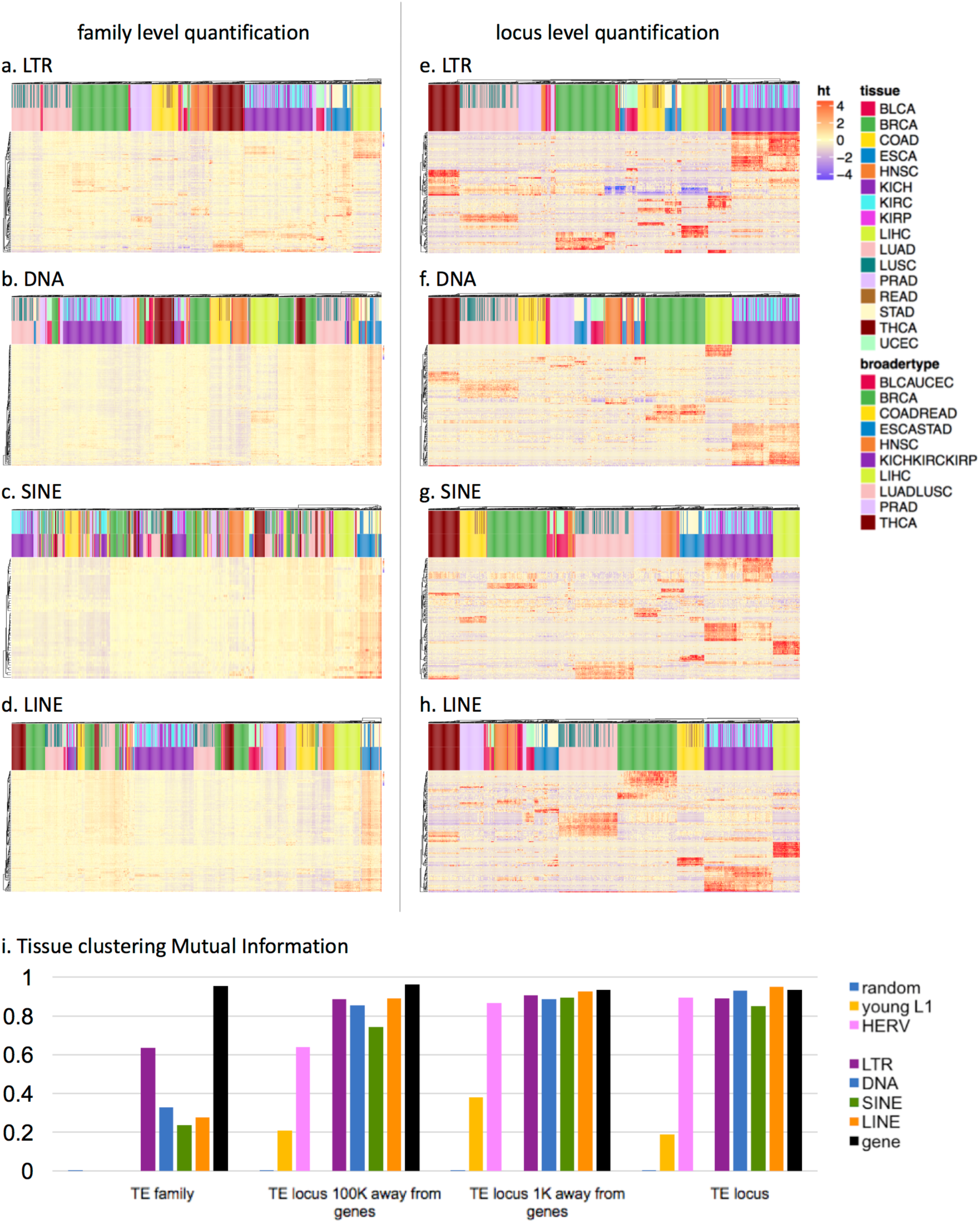
Tissue clustering based on transposable elements. Heatmap showing the tissue clustering results based on the top 150 TEs with the largest variance of each class. With the color bars at the top of the heatmap, the upper color labels show tissue types and the lower color labels show broader tissue groupings. a-d. clustering based on family level quantification. e-h. clustering based on locus level quantification. i. Clustering quality measured by Mutual Information for clustering results based on family level TE quantification, locus level quantification for TEs 100Kb away from genes, locus level quantification for TEs 1Kb away from genes, and locus level TE quantification without filtering for gene proximity.

**Figure 4.**
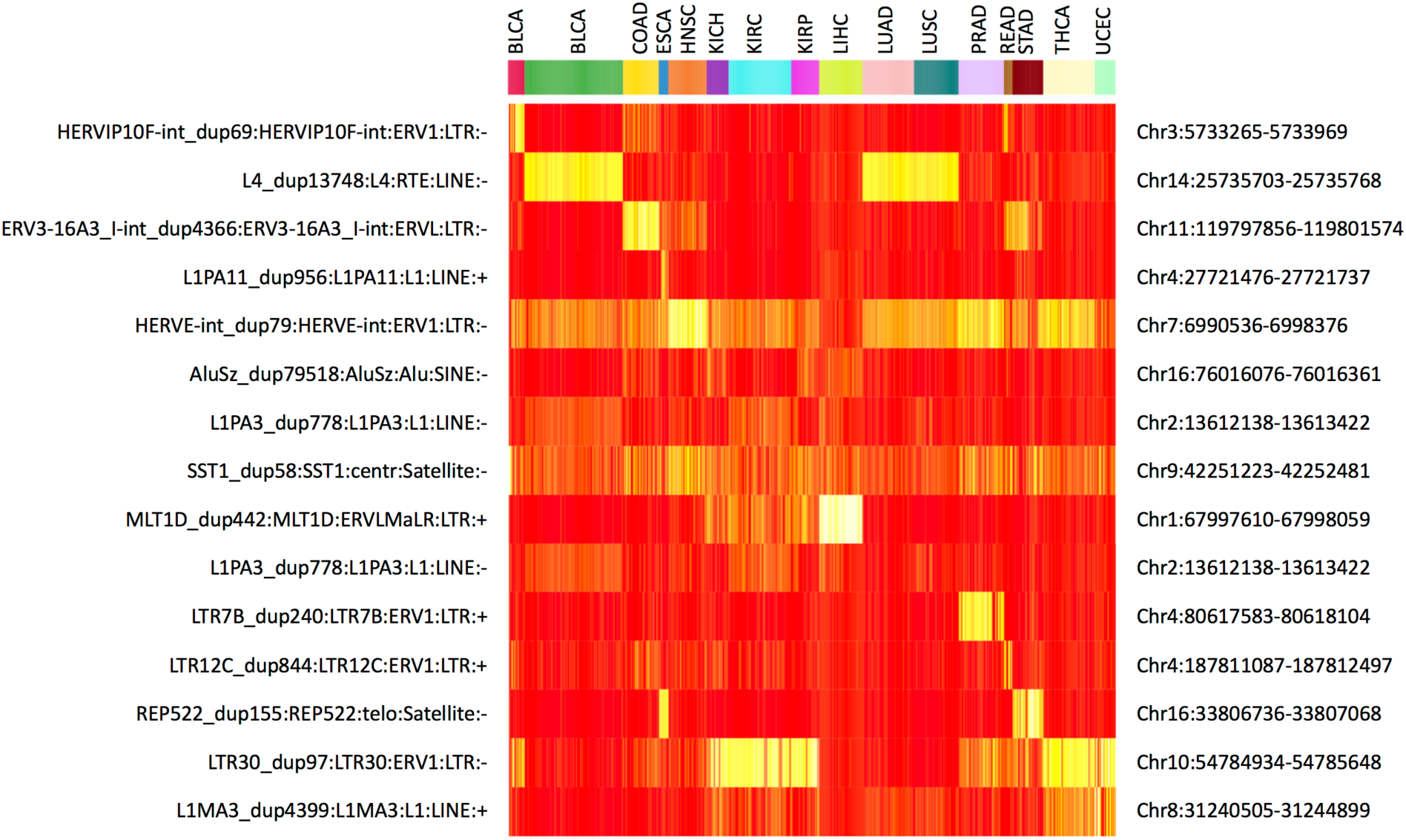
representative transposable element loci that show tissue-specific expression. 15 TE loci with tissue-specific expression was identified as representative examples among intergenic TEs that are 100Kb away from start and end of genes. Heatmap color reflects the z-score of normalized log_2_ read counts across samples.

The granular levels of locus specific TE expression contained tissue-specific information, but, the overall transcript level of TE classes did not show significant variation across tissues (Figure 5 a.). The higher expression levels for TEs seen for esophagus and stomach is confounded with the differences in sequencing protocol described above, so they are not directly comparable to the rest of the tissues. When focusing on 300 bases in the 5’ end of L1HS, it showed some variation across tissues with higher levels in the head and neck tissues and lower levels in the liver, consistent with the previous observation in adult human tissues [6] and in human cell lines [12], albeit with large within tissue variance (Figure 5 b.). Although, we cannot directly compare L1HS expression in esophagus and stomach to the rest of the other tissues, we can tell that there is clear transcription of the 5’ end of L1HS in esophagus and stomach. Figure 5 b. shows the normalized read counts in log2 scale, with a median of more than 500 reads mapping to 300 bases at the 5’ end of L1HS for esophagus and stomach (median library size 227M reads). There have been observations of full length L1HS expressed in the adult esophagus and stomach tissue, at about 80% and 150% relative to the levels in HeLa cells [6], and active L1 retrotransposition in premalignant precursor legions of esophageal adenocarcinoma [33]

**Figure 5.**
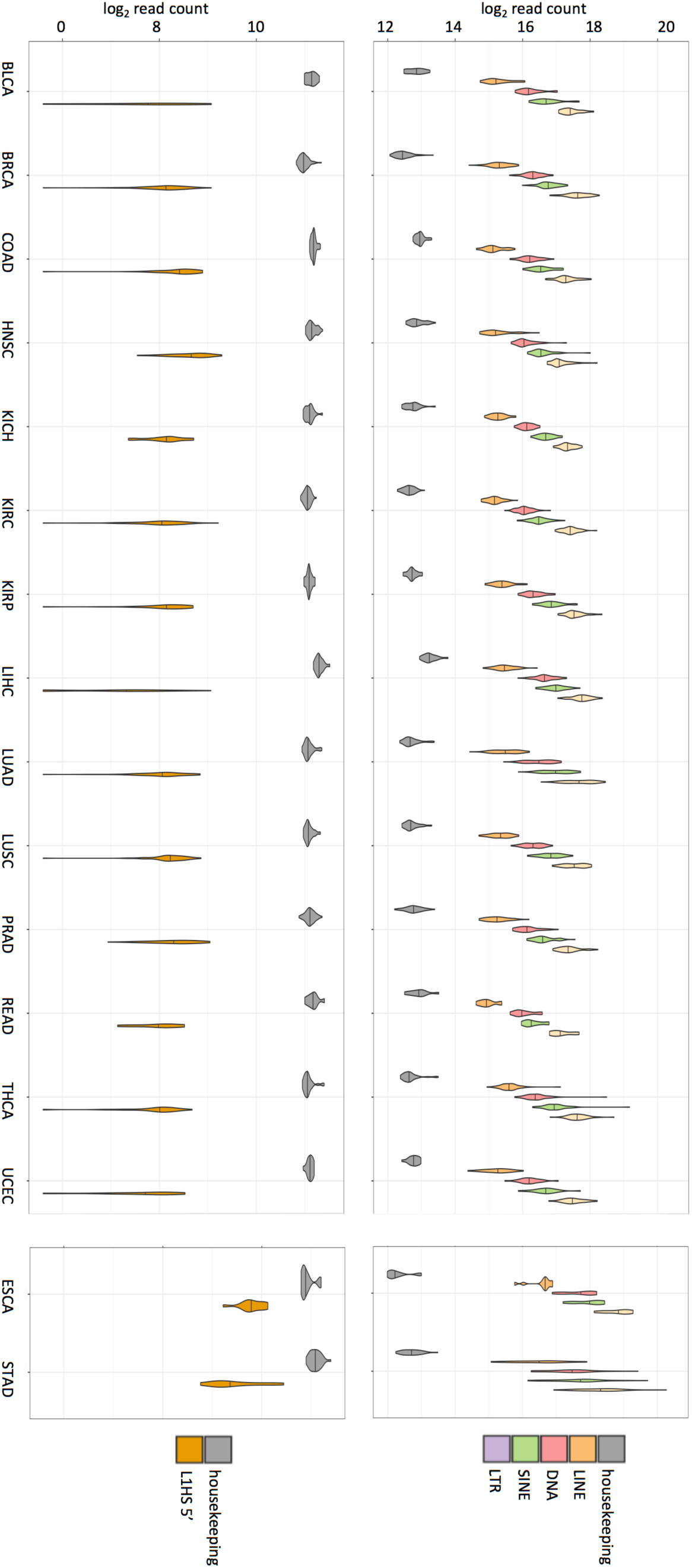
Variation in TE expression across tissues and individuals. a. Normalized and log-transformed sum of all read counts mapping to the TEs of each class, LINE, DNA, SINE and LTR are shown as violin plots. Mean read count of a set of housekeeping genes (Supplementary Table 6) are plotted as a reference. Horizontal line across the violin plot represents the median value across all samples of that tissue type. b. Normalized and log-transformed sum of all read counts mapping to 300 bp at the 5’ end of L1HS are plotted as a violin plot. Same set of housekeeping genes used in a. are plotted as a reference.

### Co-expression analysis of intergenic TEs identifies core TE modules and correlated Zinc Finger Proteins

Co-expression network analysis is an appropriate approach to examine the co-expression across different TE families and host genes together. In order to identify the common gene/TE modules that are correlated across different tissues, we did a consensus network analysis across tissues using the weighted gene co-expression network analysis in the WGCNA package [34]. For the TE family transcripts in this analysis, we only included intergenic TEs, *i.e.* we only counted reads mapping to TEs that are 1Kb away from any start and end of known genes. We identified 61 modules across 11 group of tissues, combining certain tissue types together as a broader group (colon and rectum, esophagus and stomach, kidneys, lungs). Among the 20531 genes and 992 TE families that were quantified in the 697 samples, 18670 genes and 923 TE families had enough expression level and variation to be included in the network analysis. Among those 19593 genes and TEs, 9599 genes and 658 TEs were clustered into a module of co-expression, while 9336 did not belong to any defined module. The list of modules, correlations between the modules, and topological adjacency matrix that defines the modules are visualized for the breast tissue in Supplementary Figure 4. Visualization for other tissues were similar, as we looked for consensus modules across all tissues. There were only a few modules that contained TE transcripts: only seven modules contained more than ten TE families within the module. Supplementary Table 3 shows the distribution of TE families in these seven modules. We considered modules M8, M21, M38 and M45 as core TE modules, as their membership mainly consisted of TE families as the majority (marked by * in Supplementary Figure 4).

The correlation between closely related TE subfamilies is expected because reads from transposons that map to sequences that are indistinguishable between subfamilies are assigned to multiple subfamilies with proportional weight by TEtranscripts using an Expectation-Maximization algorithm. Closely related families such as L1HS and L1PAs also share common regulatory elements at the 5’ end. But, we find that the correlated TE families in a TE module span different classes of TEs, and are replicated even when counting uniquely mapped reads only. Considering there is no sequence similarity between the SINEs, DNA transposons, LTRs and the LINEs, the correlation among these diverse class of transposons is probably due to a common regulation, or dys-regulation, that is de-repressing these transposons at the same time. There have been reports of such co-expression of ERVs and LINEs in cancerous tissues [35, 36], possibly through concordant hypomethylation [37].

There was one class of host genes that were frequently found as members of the TE co-expression modules, and they were the KRAB Zinc Finger Proteins (KZFPs). Table 2 shows the list of KZFPs that were identified as TE module members.

**Table 2.** KZFP gene members in core TE modules. KZFP genes that are members of the core TE modules, and module M3 that is correlated with core TE modules.

| core TE modules | KZFP | chromosome | KZFPs in correlated module M3 | chromosome |
| --- | --- | --- | --- | --- |
| M8 | HKR1 | 19 | KZFP169 | 9 |
|  | KZFP226 | 19 | KZFP202 | 11 |
|  | KZFP682 | 19 | KZFP266 | 19 |
|  | KZFP789 | 7 | KZFP300 | 5 |
|  | KZFP814 | 19 | KZFP320 | 19 |
| M21 | KZFP404 | 19 | KZFP431 | 19 |
|  | KZFP418 | 19 | KZFP439 | 19 |
|  | KZFP589 | 3 | KZFP44 | 19 |
|  | KZFP75A | 19 | KZFP587 | 19 |
| M38 | KZFP117 | 7 | KZFP662 | 3 |
| M45 | KZFP334 | 20 | KZFP7 | 8 |
|  | KZFP493 | 19 | KZFP700 | 19 |
|  | KZFP506 | 19 | KZFP708 | 19 |
|  | KZFP721 | 4 | KZFP714 | 19 |
|  | KZFP737 | 19 | KZFP732 | 4 |
|  |  |  | KZFP83 | 19 |
|  |  |  | KZFP841 | 19 |

### Expression of immune genes are negatively correlated with intergenic TE expression

Once we identified modules consisting mostly of transposon families, we also examined whether any co-expression modules were negatively correlated with TE modules. We found two modules, M33 and M35, that showed consistent negative correlation across tissues. The genes included in these modules were genes involved in innate immune system, interferon signaling, immunoproteasome, etc (Figure 6). Figure 6 shows the enriched annotation terms detected for both modules through the Reactome database, the top 30 genes with highest module membership for the two modules, and the correlation plot between TEs in the TE modules M8, M21, M38 and M45, and genes in module M33 and M35 in the tissues breast (Figure 6d) and esophagus/stomach (Figure 6e). Only genes and TEs that show greater than 0.6 Pearson correlation with the representative profile of the module in all tissues have been included in the correlation plot. We observe high correlation within groups and contrasting negative correlation between groups.

**Figure 6.**
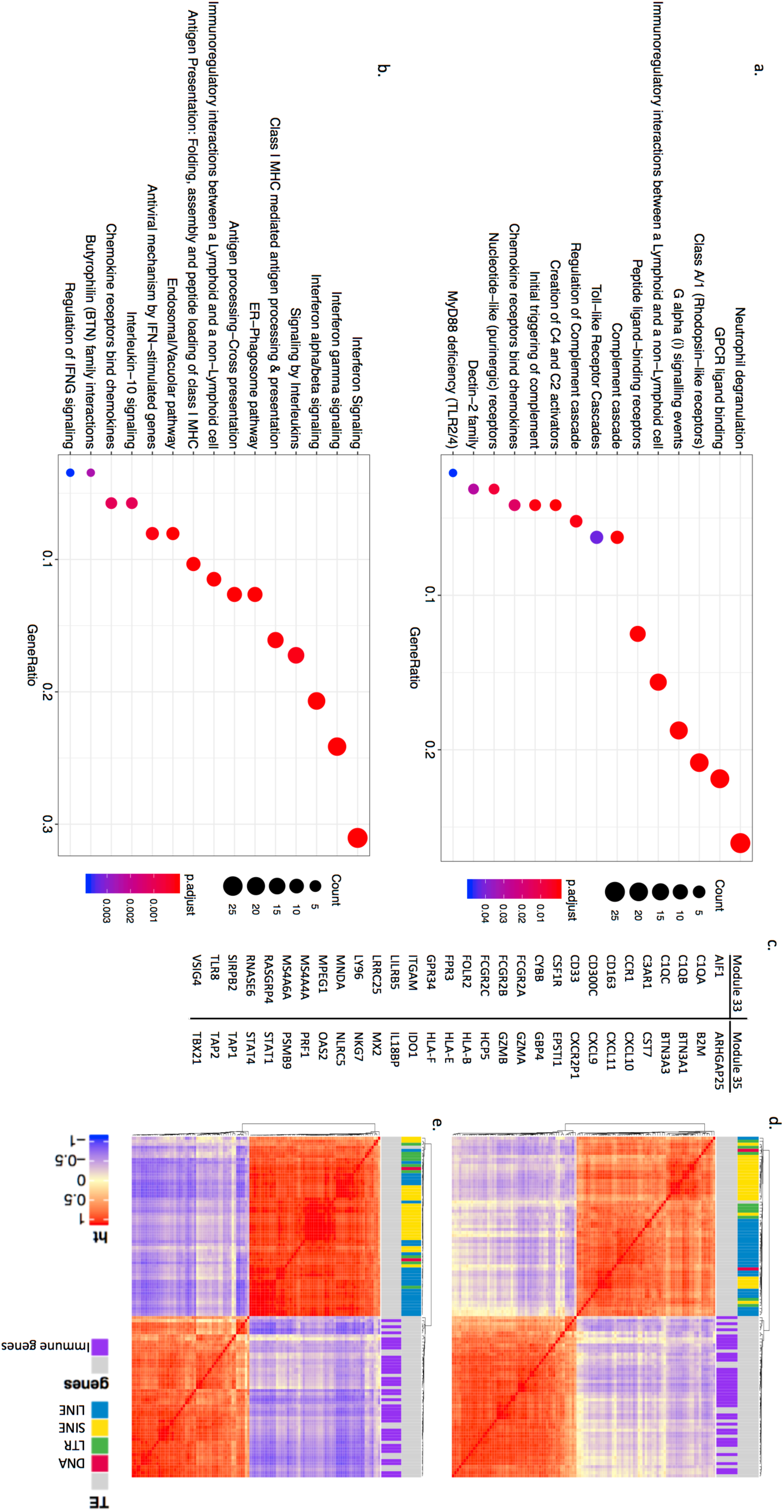
co-expression modules with immune genes are negatively correlated with TE modules. a. enriched annotations for genes belonging to module M33 identified in the Reactome database. b. enriched annotations for genes belonging to module M35 identified in the Reactome database. c. top 30 genes with highest module membership (high correlation with representative profile of the module) for each module. d-e. correlation plot showing high within group correlation and negative between group correlation between the TEs in the TE modules and the immune genes in modules M33 and M35. d. data from breast, and e. data from esophagus. Color label on top of the correlation plot show different classes of TEs, and genes that are annotated with the GO term “immune system process”.

### Co-expression analysis including intronic TEs reveals negative correlation between intronic TE expression and mitochondrial gene expression

When we include intronic TE transcripts in the overall TE expression levels, the co-expression analysis led to a different picture from the analysis of intergenic TEs. When intronic TEs are included, a single module, N1, emerges as the dominant TE module, containing 612 out of 848 TE families (72%) that was assigned a module membership. In fact, N1 consists of 72% of all TE families but only 2% of all genes.

Table 3 shows genes that are significantly correlated with module N1 in multiple tissues. A pattern immediately noticeable is that there are many pseudogenes, intronic transcripts, antisense RNAs, and long non-coding RNAs on the list. It looks like with intronic TEs, we are detecting a cell state that is dysregulated in splicing or mRNA quality control, and as a result, we are seeing a global elevation of pervasive transcription that is generally non-functional. Multiple protein coding genes on the list are involved in mRNA splicing regulation, such as *NCRNA00201*, an isoform of HNRNPU which showed strong correlation with the intronic TE module in seven different tissues, as well as *CCNL2*, *LUC7* and *LUC7L3*, perhaps as a response to the dysregulated splicing. Another interesting gene in the list is *NKTR*, hinting at the presence of immune cells in the tissue samples with high intronic TE expression. This is in contrast with the negative correlation we observe with immune genes and intergenic TE expression.

**Table 3.** Gene members in the intronic TE module N1. Genes that are members of the intronic TE module N1, the number of tissues they are assigned to module N1 in, synonyms to gene names, full gene names.

| Gene Symbol | tissues | Synonyms | Full gene name |
| --- | --- | --- | --- |
| NCRNA00201 | 7 | HNRNPU | heterogeneous nuclear ribonucleoprotein U |
| AHSA2 | 6 | AHSA2P | activator of HSP90 ATPase homolog 2, pseudogene |
| CCNL2 | 6 | CCNL2 | cyclin L2 |
| CG030 | 5 | N4BP2L2-IT2 | N4BPL2 intronic transcript 2 |
| FAM13AOS | 5 | FAM13A-AS1 | FAM13A antisense RNA 1 |
| MDM4 | 5 | MDM4 | MDM4 regulator of p53 |
| NKTR | 5 | NKTR | natural killer cell triggering receptor |
| SLC25A27 | 5 | UCP4 | solute carrier family 25 member 27 |
| ANKRD36 | 4 | ANKRD36 | ankyrin repeat domain 36 |
| LOC100190986 | 4 | LOC100190986 | uncharacterized LOC100190986 |
| LOC440944 | 4 | THUMPD3-AS1, SETD5-AS1 | THUMPD3 antisense RNA 1 |
| LOC91316 | 4 | GUSBP11 | GUSB pseudogene 11 |
| LUC7L | 4 | LUC7L | LUC7 like |
| LUC7L3 | 4 | LUC7L3 | LUC7 like 3 pre-mRNA splicing factor |
| NCRNA00105 | 4 | ASMTL-AS1 | ASMTL antisense RNA 1 |
| OGT | 4 | OGT | O-linked N-acetylglucosamine (GlcNAc) transferase |
| SEC31B | 4 | SEC31B | SEC31 homolog B, COPII coat complex component |
| KZFP789 | 4 | KZFP789 | zinc finger protein 789 |

The module that was negatively correlated with the intronic TE module (N1) included co-expression clusters consisting of mitochondrial proteins and ribosomal proteins (N4). N4 was the only module that was consistently negatively correlated with N1 with less than −0.7 correlation coefficient across all tissues. Figure 7 shows the correlation plots between TEs in N1, and genes in the mitochondrial gene module N4, for breast and esophagus. Enriched annotation terms for the genes found in the Reactome database are centered around translation and mitochondria. One intriguing possibility may be that the failed splicing and mRNA control is leading to a suppression of translation that in turn leads to reduced RNA levels of mitochondrial genes and ribosome.

**Figure 7.**
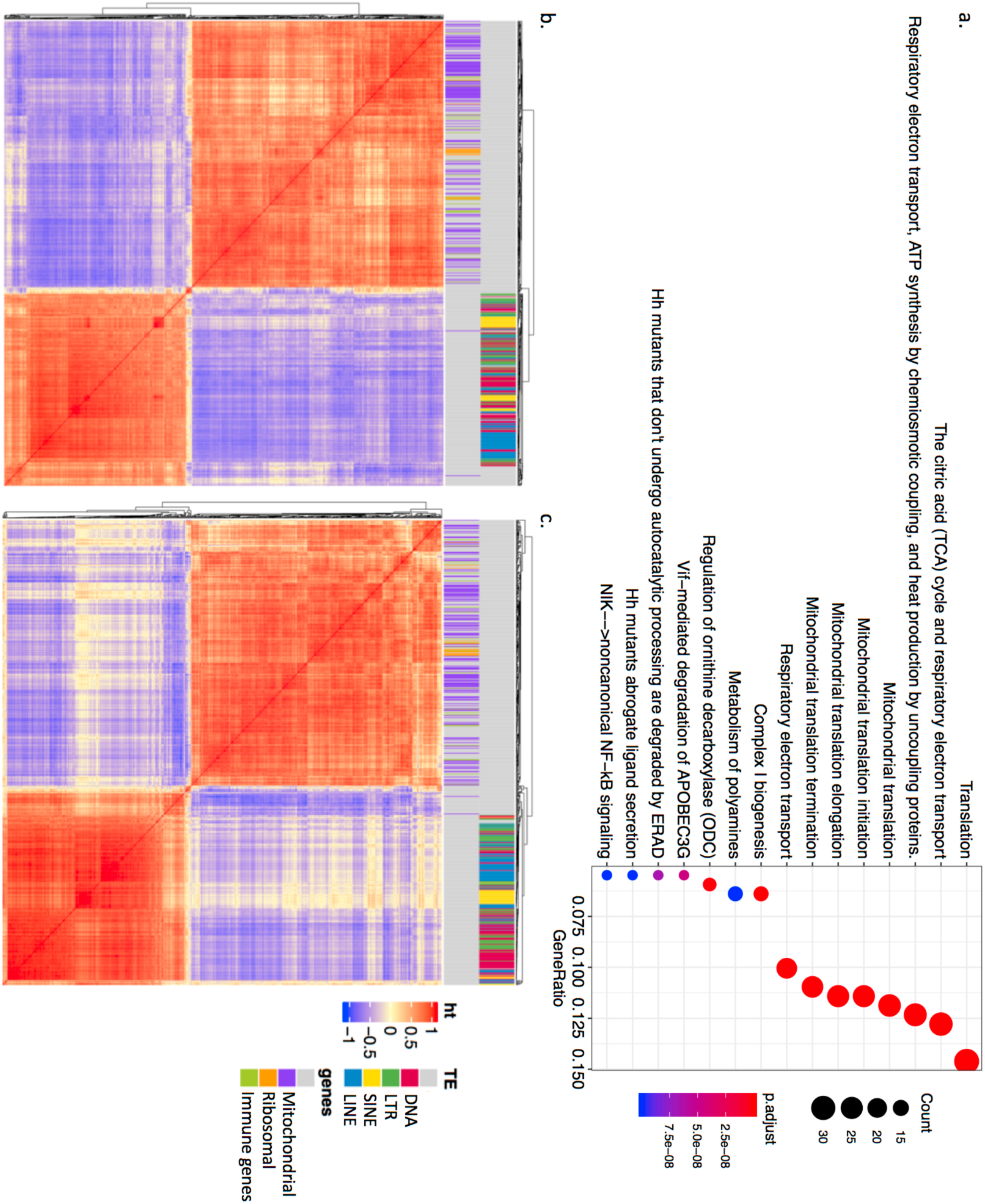
co-expression modules with mitochondrial genes and ribosome genes are negatively correlated with intronic TEs. a. enriched annotations for genes belonging to module N4 identified in the Reactome database. b-c. correlation plot showing high within group correlation and negative between group correlation between the TEs in the intronic TE module N1, and the mitochondrial and ribosomal genes in modules N4. b. shows data from breast, and c. shows data from esophagus. Color label on top of the correlation plot show different classes of TEs, and genes that are annotated with the GO term “mitochondrion” and “ribosome”.

We again saw an enrichment of KZFPs as members of the intronic TE module N1, and another module N10, that was positively correlated with N1 (Supplementary Table 4). The list of KZFPs had some overlap with the KZFPs co-expressed with intergenic TE modules, but there were some differences as well. We combined the 22 KZFPs in module N1 and N10 and examined whether there were any common transcription factor binding for these genes found in the ENCODE ChIP-seq data with the EnrichR database [38]. The region near these KZFPs were enriched with binding of *GABPA,* a regulator of nuclear encoded mitochondrial genes, in multiple cell lines (Supplementary Figure 5). This was interesting, given the negative correlation observed with intronic transposons and nuclear encoded mitochondrial gene expression described above.

### Genes co-expressed with L1HS include genes regulating major signaling pathways, chromatin, and stress response

Given the interest in the active element L1HS, and the uncertainty in L1HS quantification, we decided to limit the quantification to the 5’ region of L1HS, and examine the host genes that are specifically correlated to the expression of 5’ region of L1HS without regard to the co-expression modules. In order to control for the correlation with other TEs, especially intronic TEs, we included the representative profile of N1 as a covariate into our linear model. One concern with co-expression analysis is positional overlap. There were 14 genes that overlapped with the L1HS loci we were counting the reads from. Only 1 of the 14 genes, *RAB3GAP2*, showed significant correlation with L1HS 5’, and was removed from the final list. 56 genes were identified as negatively correlated, and 77 genes were identified as positively correlated with L1HS 5’ in at least two tissues (Figure 8, Supplementary Table 5). Notable genes include *RASA1, RASA2*, *RRAS, EGFR* and *MAPK1*, in the Ras-MAPK pathway, *ECSIT*, *TAB3* and *TRAF6*, regulators of the NF-κB pathway, *RNASEH2C*, a known L1HS repressor [39], *TET2*, known to bind to and demethylate young L1s [40], *THAP7*, a histone tail binding transcription repressor [41], and *DDI2*, a protease that cleaves and activates NFE2L1/NRF1 [42]. Multiple genes in the respiratory electron transport pathway, *ECSIT, NDUFA1, NDUFA8, NDUFB10, NDUFB8, SURF1, UQCR11, UQCRB*, were negatively correlated with L1HS 5’, even after controlling for the covariation with intronic TEs, N1. Whole list of genes are reported in Supplementary Table 5.

**Figure 8.**
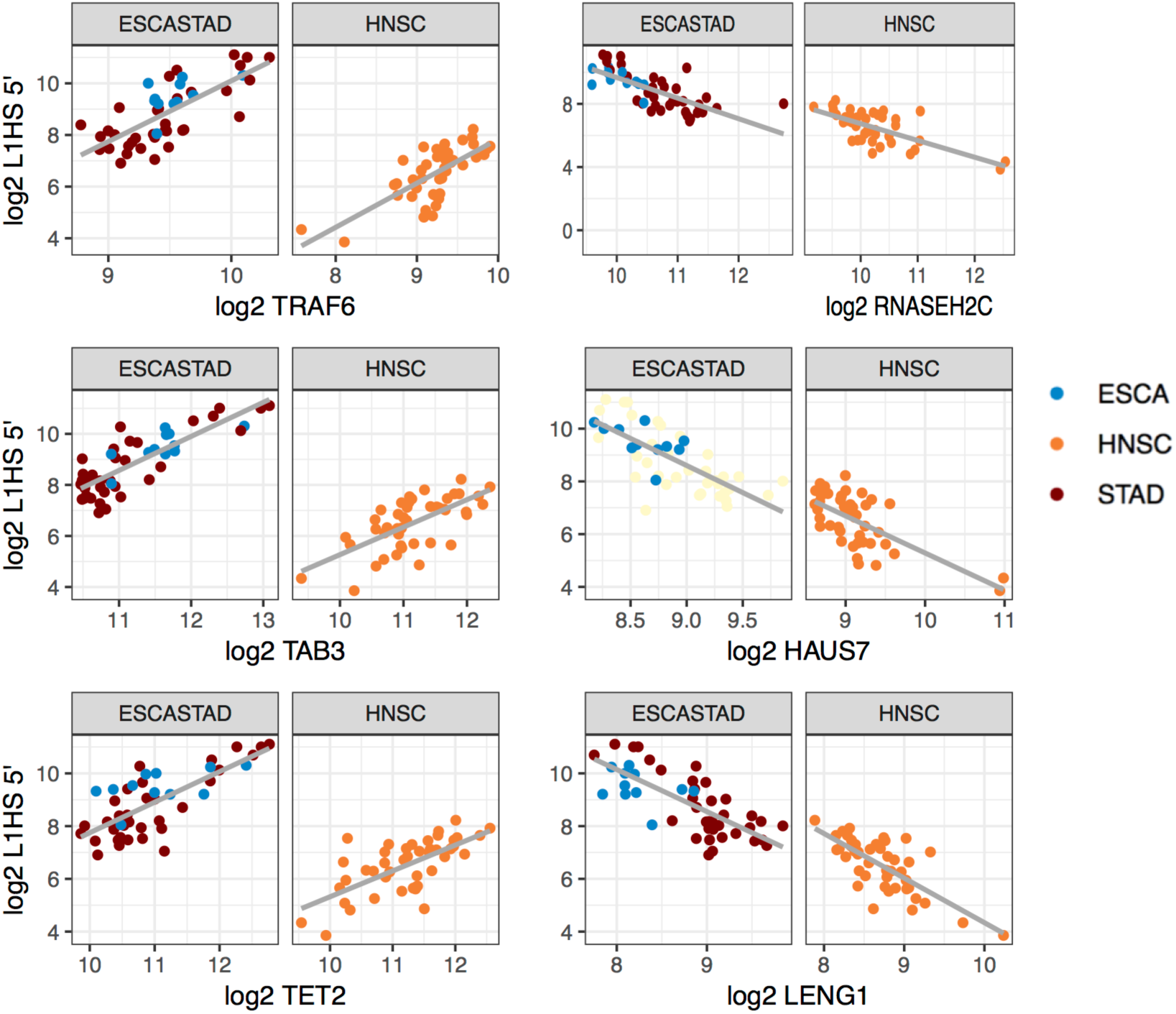
Genes that show positive and negative correlation with the transcript level of L1HS 5’ end. Gene that show significant positive and negative correlation with L1HS 5’ in multiple tissues. Esophagus and stomach are combined as one tissue group.

We checked whether the list of our negatively correlated genes were overlapping with the genes identified through CRISPR–Cas9 screen [21]. Of the 56 negatively correlated genes, three genes, *RNASEH2C*, *HAUS7*, *RNF166* were also on the list of 253 secondary screen hits. There was no overlap among the 77 positively correlated genes.

We also checked whether there were transcription factors known to bind to L1HS sequence [43] in our list. Of the 77 positively correlated genes, four genes, *YY1*, *REST*, *ELF1*, *ZBTB33* were identified to bind to L1HS [43]. There was no overlap among the 56 negatively correlated genes. To check if the same transcription factors are regulating the correlated genes and L1HS, we also checked what kind of TF binding is observed in the upstream of our correlated genes. There were a few enrichment of ENCODE transcription factor binding upstream of our list of correlated genes (Supplementary Figure 6), but except for YY1, the enriched TFs did not overlap with the list of Sun et al. [43]

### TE module expression is correlated with radiation exposure in thyroid tissue

We examined whether any of the clinical variables were associated with the TE module expression or the L1HS expression levels. We tested the variables age, days to death, pathological stage, T staging, N staging, M staging, gender, radiation and race for each tissue type. No variable was found to be associated with L1HS 5’ expression. Radiation therapy was the only clinical variable associated with module N1 (intronic TE module) expression in the non-tumorous tissue of thyroid (p-val = 0.00894, Supplementary Figure 7).

### Co-expressed TEs and KRAB-ZFPs show limited overlap with ChIP-seq binding

Based on the positive correlation observed among KZFPs and TE modules, and existing literature on the role of KZFPs for TE repression, we decided to examine the correlated expression of all pairs of 979 TE families and 366 KZFPs. The most striking pattern observed was that KZFPs and TEs show overwhelmingly positive correlation and little negative correlation. Chromosome 19, where the majority of the KZFPs are clustered, is also the chromosome with the highest density of transposable elements. This unique structure of chromosome 19 may lead to TEs embedded in KZFP genes erroneously identified as co-expressed. We avoid the confounding effect of positional overlap between TEs and KZFPs by only counting reads mapping to TEs that are in the intergenic region 1Kb away from any genes. There may be residual correlation due to shared genomic environment of a larger scale, such as the chromatin state. But, that doesn’t explain all the positive correlation, because, when we look at the locus level correlation, we find that the individual TE loci correlated with the ZNFs are scattered across all chromosomes, and not necessarily enriched on chromosome 19.

The co-expression between KZFPs and TEs were observed across almost all TE families, as 794 TE families had at least one co-expressed ZFP in at least one tissue. Certain ZFPs, such as *ZNF621*, *ZNF780B*, *ZNF84*, *ZNF33A*, and *ZNF662*, showed correlation with a wide range of TE families in multiple tissues. TE-KZFP pairs, HERVK14-int:*ZNF814*, MER57A-int:*ZNF621*, and MSTB-int:*ZNF41* were the most frequent pair-wise co-expression observed between TE families and KZFPs, found positively correlated in six different tissues. The ZFPs that were negatively correlated with TEs were *ZNF511* and *ZNF32*, but, they are not classified as KZFPs as they do not have a KRAB domain.

We looked at the family level co-expression between TE families and KZFPs and tested the overlap against the KZFP bound TE family enrichment reported in the ChIP-exo study (GSE78099 [44]). We found that there is a statistically significant association between co-expression and binding (*p-value* < 2.2e-16). But, the number of overlapping pairs were very small. Figure 9 shows the overlap between co-expression and binding enrichment. We only mark the co-expression found in at least two tissues, and we have omitted the TE-KZFP combinations that have neither co-expression nor binding enrichment from the figure. The total combinations tested that overlap between the two datasets is 200,889 (221 KZFP x 909 TE families). 4138 pairwise co-expression was observed in at least two tissues. Of those, only 119 was enriched for binding in the ChIP-exo study [44].

**Figure 9.**
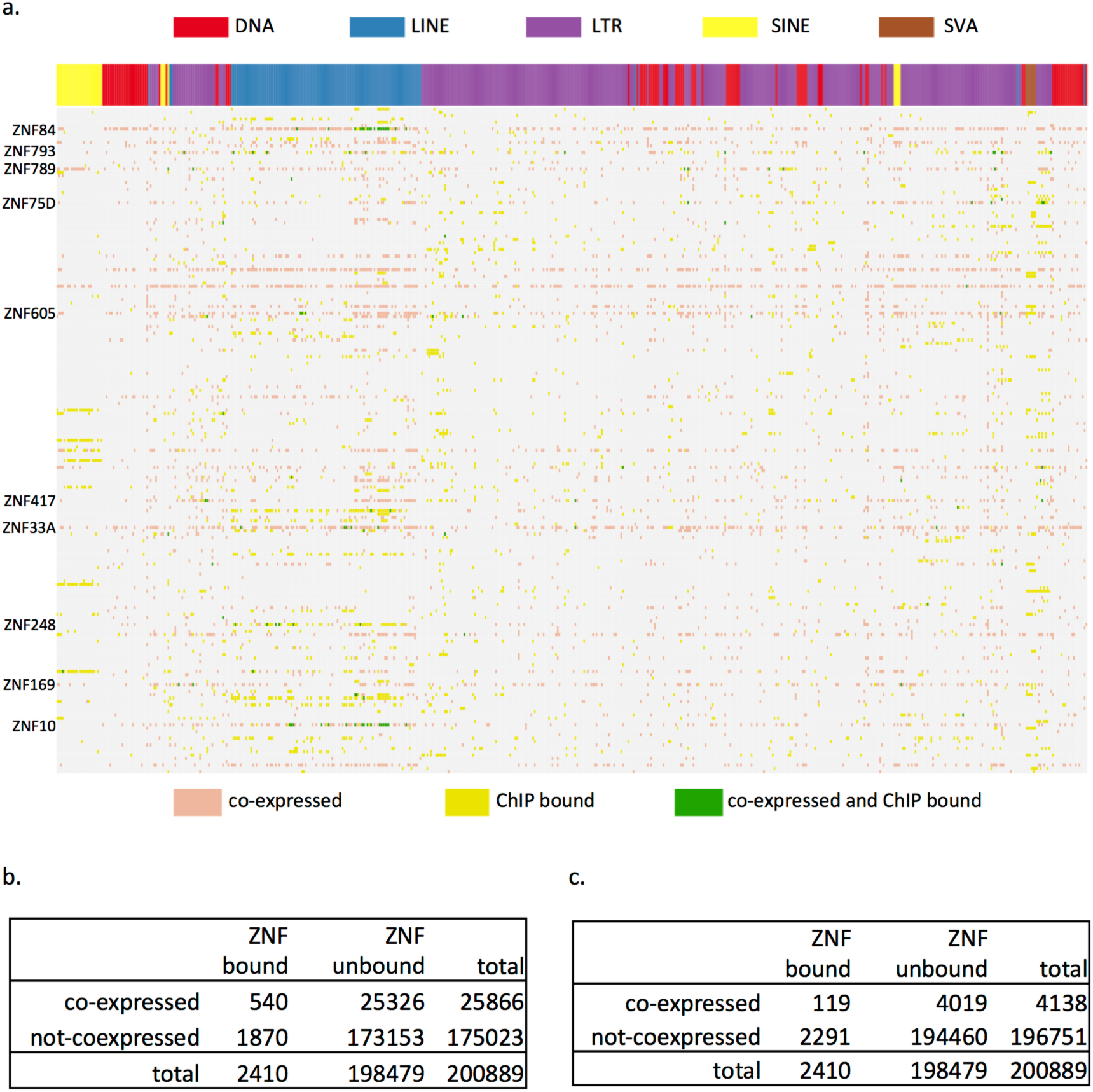
Overlap between co-expressed KZFP-TE family pairs and TE families enriched for KZFP binding. a. KZFP-TE families co-expressed in at least two tissues are marked with pink, TE families enriched for KZFP binding are marked with yellow, and TE families that are both bound by KZFP and co-expressed with same KZFP are marked with green. KZFPs that show overlap of binding and co-expression for multiple TE families are labeled along the vertical axis. b-c. categorization of co-expression and KZFP binding for all 200,889 KZFP-TE family pair-wise combinations (221 KZFP x 909 TE families), that have both expression and ChIP-exo data. b. counts co-expression significant in at least one tissue. c. counts co-expression significant in at least two tissues.

To check how the co-expression is observed at individual TE loci, we took the TE family-KZFP pairs that show correlated expression, and further tested co-expression between individual TE loci of the correlated TE family against the KZFP of interest. With correlations at the locus level, we were able to examine the locus level co-expression and compare it directly to the binding peaks reported in Imbeault et al [44]. Of the 6258 co-expressed TE loci where the KZFP had been assayed with ChIP-exo, there were only 4 that were bound by the same KZFP. We do not have a good explanation for why there is a lack of overlap between co-expression and binding at the locus level, when there was at least some amount of overlap at the family level. It looks like the co-expression we observe is a result of indirect interactions, and not necessarily direct binding.

We also observed that at the locus level, there was not a lot of overlap between the TEs that are bound by KZFPs in [44], and the TEs that are expressed in the TCGA non-tumourous tissues, regardless of the co-expression relationship with KZFPs. Here “expressed” means there is at least one sample in our data with more than five reads mapped to the TE locus, and “binding” means that there is a peak detected in the GSE78099 ChIP-seq data overlapping with the TE locus with a +-250bp buffer. Figure 10 shows the overall breakdown of the 4.5 million transposons annotated in hg19 UCSC Repeatmasker track. Statistically, there is more overlap than expected (*p-value* < 2.2e-16) between binding and expression, but, the overall proportion of TEs that are both expressed in at least one sample and bound by at least one ZFP are a tiny proportion (2.6%) of all TEs in the genome.

**Figure 10.**
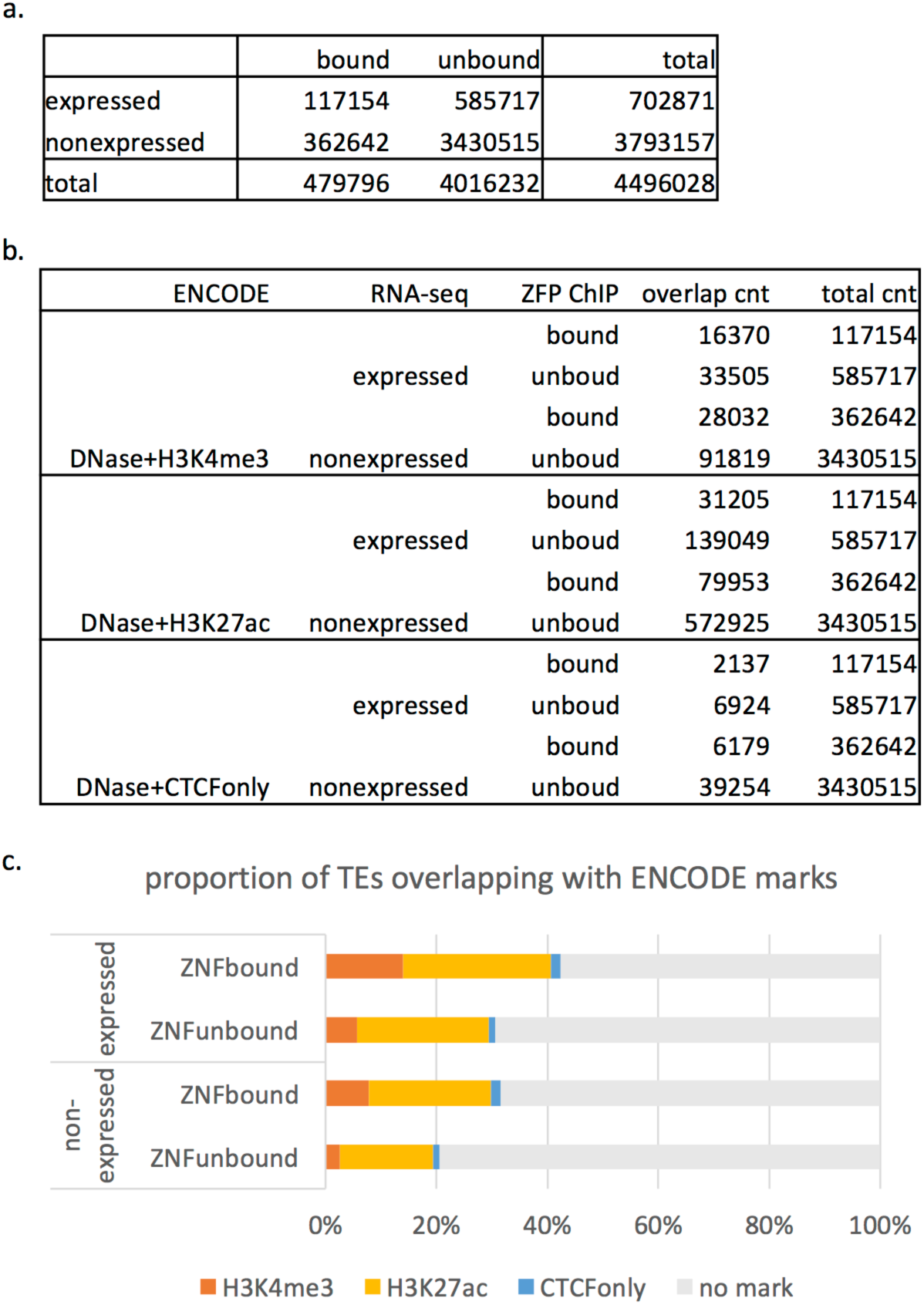
Overlap between expression and KZFP binding for TE loci. a. categorization of all 4,496,028 TE loci annotated in hg19 RepeatMasker by expression and KZFP binding. b. Overlap of expression and KZFP binding with ENCODE Candidate Regulatory Element marks. c. Proportion of each category of TEs that are marked with ENCODE Candidate Regulatory Element marks.

One interesting pattern did emerge when we examined the overlap with epigenetic marks of candidate cis-Regulatory Elements defined in the ENCODE data [45]. We were more likely to see an enhancer-like mark (DNase + H3K27ac) for TE loci that are expressed compared to non-expressed TEs, and we were more likely to see a promoter-like mark (DNase + H3K4me3) for ZFP bound TE loci compared to TEs with no binding (Figure 10). The 2.6% of TE loci that are expressed in at least one sample and bound by at least one ZFP showed the highest proportion of both promoter-like marks and enhancer-like marks. When we divide the TE loci into gene regions (genes including introns and +-1K flanking region) and intergenic regions (1K away from start and end of genes), the overall pattern remained the same, except that TE loci were twice as likely to be expressed if they are close to genes compared to intergenic regions, and the TE loci were twice as likely to be overlapping with the promoter-like marks (Supplementary Figure 8). The enhancer-like marks showed no difference between gene regions and intergenic regions, and the CTCF marks increased in the intergenic regions.

## Discussion

### Limitations to the quantification and correction

Quantifying transposon transcripts is a difficult problem, due to their ambiguity in short read mapping because of repeated content in the reference genome. Current state of the art methods rely on Expectation-Maximization to account for the uncertainty in multi-mapped reads [29]. Focusing only on uniquely mapped reads doesn’t really solve this problem, and will lead to biased quantification, favoring older elements with higher mappability. Scott et al. have demonstrated that by relying on unique mutations found within individual L1HS loci, and by including sequences of non-reference polymorphic L1HS loci, it is possible to identify the source of the L1HS activity with substantial success [46]. But, in our study we did not attempt to identify the individual loci of L1HS transcription, and instead focused on the totality of reads mapping to regions of annotated L1HS that align to the 5’ end of L1HS consensus sequence.

Another complication in TE transcript quantification is that TEs are frequently embedded within introns that are transcribed before they are processed, or sometimes fail to be spliced out, or embedded within exons or non-coding RNAs that are expressed in different conditions [28]. To account for this source of error, we introduced a method to correct for TE reads coming from retained introns or pre-mRNA. Although we observed large corrections for specific transposable elements embedded within introns, the correction is not complete. We can tell this from the observation that the co-expression profiles of intronic TEs are different from the co-expression profiles of intergenic TEs away from the genes. The genes co-expressed with intronic TEs include pseudogenes, intronic transcripts, anti-sense transcripts and genes with functions in splicing. A more accurate approach would be to correct for the read counts from retained introns before the EM algorithm based on the read depth of uniquely mapped reads, and then run EM based on the corrected counts. But, estimating the read depth of the repeat region using uniquely mapped reads is a difficult problem. The effective length of the uniquely mapped region is difficult to estimate, because again mappability varies from locus to locus for any TE, depending on the unique mutations it has accumulated. So, for this study, we decided to use the easier approach to run EM first, and probabilistically assign the TE reads, and then correct based on the expected read depth across the length of the TE locus. An important future study would be to study the mappability of individual TE loci carefully, including the known polymorphic sites, and to design a software for TE quantification that can take into account the mappability of each locus in its EM algorithm, as well as correct for the retained introns while considering the effective length of the uniquely mappable region within the TE.

Despite these limits, the main results of co-expression analysis were not affected by the quantification. Most of the results in the paper were replicated when quantification was done on uniquely mapped reads only. The only results that changed between the multi-mapped approach vs. the unique mapping approach were the genes correlated with the L1HS 5’ expression level. For those, we decided to report on results from the multi-mapped reads rather than the unique reads, because of the bias of the uniquely mapped reads we described above.

### Stress, immune response and TE expression

Initially, when we started the project, our goal was to identify candidate genes involved in transposon control, based on the co-expression analysis. But, once the analysis was done, the results were pointing to what induces TE expression, rather than what suppresses TE expression. Among the genes known to function in transposon control, *RNaseH2C* (Figure 8), *HAUS7*, and *RNF166* [21] showed negative correlation with L1HS. But several well-known genes with functions in transposon control, *e.g. MORC2, SIRT6, KAP1, SAMHD1, MOV10, ZAP, C12orf35* (human ortholog of *RESF1*), etc. are missing in our list of significantly correlated transcripts. Instead, the major theme that emerged from our results is signal transduction, immune response, and stress response as seen in the correlation between L1HS and *DDI2*, Ras-MAPK and NF-κB pathway. In humans, various stresses have been shown to induce LINE1 transcription or activation including chemical compounds [47–49], radiation [50, 51], oxidative stress [52] and aging [53]. Most of these studies have observed L1 activity *in vitro*, by exposing cultured cells to stress factors and assaying the retrotransposition activity.

The negative correlation we find between TE expression and immune gene activity has been reported before in gastrointestinal cancer samples. Jung et al. have shown that the L1 retrotransposition rate is inversely correlated with expression of immunologic response genes [54]. Here, we extend those results and show that the negative correlation between TE expression and immune response is a pattern found in non-tumorous samples as well, across different tissues and different classes of TEs. This relationship is confusing, since it is opposite of the positive correlation we find between L1HS and NF-κB pathway genes (Figure 8), and opposite of the pattern observed in several cancer studies, where DNA hypomethylation and expression of endogeneous retrovirus activates interferon signaling [55–57]. Immune active environment surrounding these tumor adjacent cells plus nucleic acids in the extra-cellular environment coming from cancer nearby may be putting the tumor adjacent cells in an antiviral state. It is known that interferon signaling induces proteins that act against viruses. *ZAP* is one example that degrades viral RNA as well as RNA of LINEs and Alus [58], although *ZAP* does not show correlated expression with the TE modules in our data. We hypothesize that such cell states may reduce transposon transcripts with higher sensitivity through RNA degradation and chromatin remodeling.

The tissue samples in this study are not representative of “normal” cells, as they are collected as controls from tissue adjacent to cancer cells. Although they are not undergoing the molecular changes associated with malignant transformation, they could be under the influence of nearby environment, with changes in pH levels, inflammation, and infiltration of immune cells. The inclusion criteria for TCGA does not allow patients with any prior systemic chemotherapy or any other neoadjuvant therapy, but it does allow local radiation, and we observe that past local radiation is associated with higher TE expression levels in adjacent cells in thyroid tissues. Given the characteristics of the samples, the variation in TE expression levels or the co-expression pattern we observe in this study may be due to cancer-associated stress. Future studies will be needed to confirm whether the results are replicated in true normal tissue.

### TEs and KRAB-ZFPs

ChIP-Seq studies on KRAB-ZFPs have identified extensive binding between this family of proteins and transposable elements [44, 59], implying a role for suppressing TE expression. KRAB domain is a well-known repressor domain and together with the co-factor KAP1 (TRIM28), the KZFP-KAP1 complex has been shown to silence both exogenous retroviruses and endogenous retroelements during embryonic development [60, 61]. Based on this observation, and the pattern of co-evolution of retroviral LTRs and the C2H2-Zinc Finger gene family, it has been hypothesized that the KRAB-ZFPs function in transposable element suppression [62]. But except for a few KRAB-ZFPs, most members do not have a characterized function. In an alternative hypothesis, instead of its original role in silencing, it was proposed that KRAB-ZFPs may also have a role in controlling domesticated transposable elements that contribute to the host transcription regulation network [63]. In our co-expression analysis, we found overwhelming positive correlation between KZFPs and TEs across all classes of TEs. This positive correlation was observed whether we are counting multi-mapped reads or uniquely mapped reads, and whether we are counting TEs close to genes, or TEs in the intergenic regions. Despite this robust positive correlation, we found that the co-expressed relationship showed limited correspondence with published ChIP-seq binding results. There was statistically meaningful but very small number of overlap at the family level, and almost no overlap at the locus level. The co-expression we observe seems to be largely an indirect relationship, and not a result of direct binding. There have been observations of a unique chromatin state that is shared between ZFP clusters and repeat classes by the Roadmap Epigenomics project [64]. This chromatin state, termed ZNF/Rpts, is characterized by H3K36me3 marks co-occuring with H3K9me3 marks and high DNA methylation. It is possible that local chromatin environment that is co-regulated at a larger scale is responsible for the correlation at the RNA level.

## Conclusions

TE derived transcripts in the non-tumourous tissues show large variation across tissues, and across individuals. Co-expression network analysis within tissues revealed general co-expression of TEs across all classes. It also found strong co-expression between TEs and KRAB-Zinc Finger Proteins that are replicated in multiple tissues, but not congruent with direct binding of TE-ZFP relationships assayed through ChIP-seq. We also found negative correlation between intronic TEs and mitochondrial genes, and between intergenic TEs and immune response genes, replicated in multiple tissues.

## Methods

### RNA-Seq and gene expression quantification in the non-tumorous tissues

We used the gene level quantification provided by The Cancer Genome Atlas (TCGA) for the gene expressions [24–26]. We collected gene level quantifications for 697 samples from TCGA. We focused on cancer types that had at least 10 control samples of RNA-seq data, collected from non-tumorous tissue adjacent to the cancer tissue. As a result, 16 different tissue types were included in our analysis: BLCA (Bladder urothelial carcinoma), BRCA (Breast carcinoma), COAD (Colon adenocarcinoma), ESCA (Esophageal adenocarcinoma), HNSC (Head and neck squamous cell carcinoma), KICH (kidney chromophobe), KIRC (kidney renal clear cell carcinoma), KIRP (Kidney renal papillary cell carcinoma), LIHC (Liver hepatocellular carcinoma), LUAD (Lung adenocarcinoma), LUSC (Lung squamous cell carcinoma), PRAD (Prostate adenocarcinoma), READ (Rectum adenocarcinoma), STAD (Stomach adenocarcinoma), THCA (Thyroid carcinoma) and UCEC (Uterine Corpus Endometrial Carcinoma). Number of samples for each tissue is described in Supplementary Table 1. Although we will use the acronym for the cancer type to describe these tissues, we emphasize again that all our samples come from the non-tumorous tissues collected from the same organ of the same patient with the cancer. The cancer tissue samples were not included in our analysis.

Methods for sequencing and data processing of RNA using the RNA-seq protocol for all tissues except esophagus and stomach have been previously described for TCGA in [24–26]. Briefly, RNA was extracted, prepared into poly(A) enriched Illumina TruSeq mRNA libraries, sequenced by Illumina HiSeq2000 (resulting in paired 48-nt reads), and subjected to quality control. Sequencing for esophagus and stomach was done differently from other tissues and have been described in [27]. Briefly, poly A+ mRNA was purified using MultiMACS mRNA isolation kit on MultiMACS 96 separator, and double stranded cDNA was synthesized using the Superscript Double-Stranded cDNA synthesis kit. Following the library preparation protocol described in [27], the final DNA was sequenced on Illumina HiSeq2000 with paired end 75-nt reads. RNA reads were aligned to the hg19 genome assembly using Mapsplice [65]. Gene expression was quantified for the transcript models corresponding to the TCGA GAF2.1 using RSEM [31]. We used the raw_count values in the .rsem.genes.results files, rounded to an integer, as the gene level quantification.

### Quantifying TE derived transcripts at the locus and family level

We collected RNA-seq level 1 binary alignment files (.bam files) for 697 samples (Supplementary Table 1) from TCGA. The bam files were then converted to fastq and realigned to the hg19 reference genome using STAR and Bowtie1. With the STAR alignment, we allowed up to 200 mappings for every read (--outFilterMultimapNmax 200 --winAnchorMultimapNmax 200). With the Bowtie1 alignment, we only allowed the single best alignment for each read, and if there were multiple best alignments, the read was discarded from the final alignment (-m 1 -S -y -v 3 -X 1000 --max). We used a modified version of the software TEtranscripts [29] for quantifying the reads mapping to annotated transposons. TEtranscripts is a software that can quantify both gene and TE transcript levels from RNAseq experiments. It takes into account the ambiguously mapped TE-associated reads by proportionally assigning read counts to the corresponding TE families using an Expectation-Maximization algorithm. We implemented two modification to the original TEtranscripts software. 1) We modified it to report read counts for each individual TE locus in the reference genome in addition to the family level counts. 2) We developed a function to discount the read counts by removing read counts that correspond to transcripts containing TE sequences that originate from pre-mRNA or retained introns in the mature RNA [28]. Downstream analyses were done using the discounted quantification based on multi-mapped reads and the uniquely mapped quantification for both the STAR alignment and the Bowtie1 alignment, to assess the impact of uncertainty in multi-mapped reads.

The retrotransposon annotations used were generated from the RepeatMasker tables, obtained from the UCSC genome database and provided by TEtranscripts. For quantifying reads mapping to the TE flanking introns we generated gtf files containing 1) the TE flanking intron positions, 2) the intergenic TE positions, and 3) the exonic TE positions (TEs that fall within an exon, including non-coding RNA genes). In case of intronic TEs, we use the algorithm described above to discount the transcripts from pre-mRNA or retained introns. In case of intergenic TEs, we count all EM estimated reads mapped to TEs without any discount. In case of exonic TEs, we ignore those counts altogether, and the exonic TEs do not contribute to the locus count nor the family level count.

### Normalization and transformation of read counts

After quantifying the reads mapping to annotated genes and TEs, both the gene level counts, and the TE counts were normalized between samples across all tissue types with DEseq2. We used the default “median ratio method” for normalization in DESeq2 [66]. Briefly, the scaling factor for each sample is calculated as median of the ratio, for each gene, of its read count to its geometric mean across all samples. The assumption of the median ratio method is that most genes are not consistently differentially expressed between tissues. If there is systematic difference in ratio between samples, the median ratio will capture the size relationship. But, this assumption may be violated when we are comparing large number of tissues types at the same time, since a large proportion of the genes may be differentially expressed in at least one tissue type, or one of the tissues may be extremely biased in their number of differentially expressed genes. In order to achieve more robust normalization, we used a two-step normalization method called the differentially expressed genes elimination strategy (DEGES) [67]. We performed preliminary normalization using the “median ratio method”, filtered out potential differentially expressed genes in the data, found a subset of robust non-differentially expressed genes, and used the subset to perform the second round of “median ratio normalization”. The resulting pairwise MA plot between tissues after normalization showed better normalization compared to the regular one-step normalization. The size factors for each sample obtained from the two-step normalization on gene counts were then used to normalize the TE quantifications of the same sample. The normalized counts were log2 transformed using the variance stabilizing transformation function in DESeq2 [66, 68] for downstream analysis.

### Clustering of samples by expression pattern

We cluster the samples using the “average” method (=UPGMA) in the *hclust* function of R, and visualize the clusters with the ComplexHeatmap package [69]. The top 150 genes or TEs, with the largest variances on the log2 transformed read counts were used for clustering. We did not select genes by any measure of differential expression across tissues. These genes were simply the genes showing the largest variance in read count across all 697 samples, regardless of tissue type. We exclude genes and TEs on X and Y chromosomes. Based on the log2 read count of the top 150 TEs, a dissimilarity matrix is calculated and used for the clustering and visualization. The average method of *hclust* computes all pairwise dissimilarities between the members of the two clusters and considers the average as the distance between the two clusters. Hierarchical clustering starts with each sample assigned to its own cluster and then proceeds iteratively, at each stage joining the two most similar clusters, continuing until there is just a single cluster. For the locus level TE expression, we filtered out all loci that had less than 5 read counts for every sample. To compare the clustering of samples based on gene expression, TE expression and random assignment, we used the normalized Mutual Information (NMI) measure [70]. The hierarchical clusters were cut off at k = 16, the number of different tissue types. Because the resulting clusters were not accurate enough to distinguish between similar tissues, we used a broader tissue grouping to compare with the clusters. The tissues were grouped to 10 broader types based on preliminary clustering: bladder/endometrium (BLCA, UCEC), breast (BRCA), liver (LIHC), colon/rectum (COAD, READ), esophagus/stomach (ESCA, STAD), head and neck — the squamous epithelium in the mucosal surfaces inside the mouth, nose, and throat (HNSC), kidney (KICH, KIRC, KIRP), lung (LUAD, LUSC), prostate (PRAD), and thyroid (THCA). The broader tissue type of each sample was used as the ground truth. Each resulting cluster was then assigned a group label based on the majority tissue type. Normalized mutual information was calculated by comparing the labels from the clustering to the true class labels. Random assignment clusters were generated by permuting the tissue types with and without replacement 100 times, and the mean NMI was reported.

### Co-expression network analysis with TEs and host genes

Weighted correlation network analysis was done with the WGCNA package [34]. We start with the signed pair-wise correlation matrix across the expression levels (normalized log_2_ read counts) of all genes and TE families. We calculate the adjacency matrix by raising the correlation matrix to the power of 14, power parameter selected using the scale free topology measure, effectively suppressing the low correlations due to noise. Topological overlap based distance matrix (TOM) is calculated using the network topology resulting from the adjacency matrix. This procedure was repeated for each tissue, and a consensus TOM was calculated across all tissues. We used hierarchical clustering on this consensus topological overlap matrix to identify clusters (modules) that are shared across tissues. A representative gene expression profile of the module is defined by the first principal component of the expression levels of all members in each module. The representative profile is compared between each module to identify positive and negative correlation between modules.

### Correlated expression between genes and L1HS 5’

We blasted all the L1HS instances annotated in repeatmasker against the L1HS consensus sequence and identified the regions aligning to the 300 bases of the 5’ end of the consensus sequence. We counted all the reads mapping to the list of L1HS 5’ ends and normalized them with the same size factor described above. We used log2 transformed value of this normalized read count as the variable representing L1HS transcript level. Correlation between gene and L1HS 5’ transcripts were tested in each tissue groups separately, in bladder, breast, liver, colon/rectum, stomach/esophagus, head and neck, kidney, lung, prostate and thyroid. We tested 20532 genes for each tissue group using a linear model with log2 L1HS 5’ expression as the dependent variable, and log2 gene expression as the independent variable. For a gene to be included in our test, it had to be present in at least eight individual patients. We also required that the gene be expressed with a minimum RPM of 2 in 75% of the samples to be included in the dataset. In addition to the radiation therapy for thyroid tissue, we considered effective library size (sum of all normalized counts) and the batch ID provided by the TCGA project as additional covariates. Since there was significant co-expression across all TE classes especially for the intronic TEs, we included the expression profile of the intronic TE module N1 identified during the co-expression network analysis as a covariate in our linear model. The linear model we used is described below.

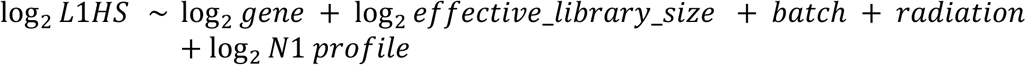

We tested all combination of linear models that can be created by including or excluding these variables. Second-order Akaike Information Criterion (AIC_c_) was used to select the best linear model. We used the coefficient and p-value from the best model to calculate the q-values. Genes with q-value < 0.0001 in at least two tissues were identified as correlated genes.

### Correlation between TE and KRAB-ZFPs

To understand the positive correlation between TEs and KRAB-ZFPs, we looked at the correlation between each KZFPs and TEs at the family level and at the individual TE locus level in different tissue types. We tested the correlation for 366 KRAB Zinc Finger Proteins that were identified in Imbeault et al. [44] and also found in our gene expression data. Because the search space of pairwise combinations of KZFP and individual TE loci was too large, we examined the relationship in a step-wise approach. In the first step, we tested the correlation between all pairwise combinations of 366 KZFPs and 979 TE subfamilies using the TE quantification at the family level in each tissue type. Then, in the second step, once the significantly correlated KZFP and TE family was identified, we focused on those pairs. We tested the correlation between the expression of the significant KZFP and the expression of each individual locus of the significant TE family in the tissue where the initial co-expression was found to identify individual TE loci that are co-expressed with the KZFP.

Overlap between co-expression and binding was examined at the family level and at the locus level. At the family level, we downloaded the family enrichment results from Imbeault et al. [44] and identified pairs of TE families and KZFP that had an enrichment score greater than 1. We compared those families enriched with binding of specific KZFPs to our co-expression results, to check if the TE families were co-expressed with the same KZFPs. At the locus level, we compared the co-expressed TE loci with the binding peaks reported in the dataset GSE78099. We took +- 250 bp around the boundary of peaks and found overlap with TE annotations from Repeatmasker. We checked if the TE locus overlapping with ChIP-seq peaks were found to be co-expressed with any KZFPs.

## List of Abbreviations

TE: Transposable Element
KZFP: KRAB Zinc Finger Protein
TCGA: The Cancer Genome Atlas

## Declaration

### Ethics approval and consent to participate

Not applicable.

### Consent for publication

Not applicable

### Availability of data and material

The datasets generated and/or analysed during the current study are available in the github repository, https://github.com/HanLabUNLV/TEcoex. The modified version of the TEtranscripts software [29] and the required gtf files can be found at https://github.com/HanLabUNLV/tetoolkit.

## Authors’ contributions

NC performed the co-expression analysis. GMJ performed TEtranscripts quantification. SQ performed the REC score analysis. NC, AR analyzed the KZFP-TE locus pairwise co-expression and KZFP motif search. CS re-ran the pipeline with the Bowtie and STAR alignment program. AA, CC, DC, ON assisted with the analyses. MVH designed the experiments, modified the TEtranscripts software, analyzed and interpreted the data. MVH wrote the manuscript with the help of all other authors. All authors read and approved the final manuscript.

## Acknowledgements

We thank the TCGA and GTEx project teams for making the data available.

