## Supplementary Materials for "Transcriptome analyses of tumor-adjacent somatic tissues reveal genes co-expressed with transposable elements"

### Additional Files

#### Supplementary Table 1. Tissue types and the number of normal tissue samples

RNA-seq data of non-tumorous samples were collected from the Cancer Genome Atlas.

BLCA (Bladder urothelial carcinoma) , BRCA (Breast carcinoma), COAD (Colon adenocarcinoma), ESCA (Esophageal adenocarcinoma), HNSC (Head and neck squamous cell carcinoma), KICH (kidney chromophobe ), KIRC (kidney renal clear cell carcinoma ), KIRP (Kidney renal papillary cell carcinoma ), LIHC (Liver hepatocellular carcinoma), LUAD (Lung adenocarcinoma), LUSC (Lung squamous cell carcinoma), PRAD (Prostate adenocarcinoma), READ (Rectum adenocarcinoma), STAD (Stomach adenocarcinoma), THCA (Thyroid carcinoma) and UCEC (Uterine Corpus Endometrial Carcinoma)

| tissue | samples |
|---|---|
| BLCA | 19 |
| BRCA | 113 |
| COAD | 41 |
| ESCA | 11 |
| HNSC | 44 |
| KICH | 25 |
| KIRC | 72 |
| KIRP | 32 |
| LIHC | 50 |
| LUAD | 59 |
| LUSC | 51 |
| PRAD | 52 |
| READ | 10 |
| STAD | 35 |
| THCA | 59 |
| UCEC | 24 |
| Total | 697 |

#### Supplementary Table 2. Tissue clustering results based on TE expression

Normalized Mutual Information values for tissue clustering, based on different classes of transposable element expression.

| | TE family | TE locus 100K away from genes | TE locus 1K away from genes | TE locus |
|---|---|---|---|---|
| random | 3.13E-04 | 2.19E-04 | 2.75E-04 | 2.38E-04 |
| young L1 | | 0.209 | 0.381 | 0.189 |
| HERV | | 0.639 | 0.867 | 0.895 |
| LTR | 0.633 | 0.887 | 0.907 | 0.890 |
| DNA | 0.326 | 0.852 | 0.887 | 0.929 |
| SINE | 0.238 | 0.742 | 0.892 | 0.851 |
| LINE | 0.275 | 0.890 | 0.927 | 0.950 |
| gene | 0.955 | 0.962 | 0.935 | 0.935 |

#### Supplementary Table 3. TE modules and TE family memberships

TE co-oexpression modules and the TE families belonging to each TE module. We consider the top four modules, M8, M21, M38, M45  with the largest TE family membership as core TE modules. M3 is not defined as a TE module, since it has more gene members than TE family members, but it showed strong correlation with all four TE modules.

| Module | total size | TE family | Alu | CR1 | DNA | ERV1 | ERVK | ERVL | MaLR | Gypsy | Blackjack | Charlie | Tip100 | L1 | L2 | MIR | MuDR | PiggyBac | Satellite | TcMar | Mariner | Tc2 | Tigger | UCON |
|---|---|---|---|---|---|---|---|---|---|---|---|---|---|---|---|---|---|---|---|---|---|---|---|---|
| M8 | 285 | 204 | 23 | 0 | 0 | 65 | 11 | 20 | 16 | 2 | 3 | 8 | 4 | 29 | 0 | 0 | 1 | 0 | 1 | 0 | 1 | 3 | 9 | 1 |
| M21 | 182 | 137 | 3 | 3 | 1 | 25 | 2 | 16 | 15 | 4 | 2 | 18 | 5 | 27 | 0 | 0 | 1 | 2 | 0 | 0 | 2 | 0 | 7 | 0 |
| M38 | 122 | 91 | 3 | 0 | 0 | 22 | 1 | 5 | 9 | 2 | 1 | 6 | 0 | 29 | 0 | 0 | 0 | 0 | 2 | 0 | 0 | 1 | 7 | 0 |
| M45 | 107 | 70 | 3 | 0 | 0 | 11 | 2 | 9 | 6 | 0 | 0 | 5 | 4 | 14 | 4 | 4 | 0 | 0 | 0 | 1 | 1 | 0 | 3 | 0 |
| M3 | 341 | 53 | 0 | 1 | 1 | 23 | 3 | 8 | 4 | 1 | 1 | 2 | 1 | 2 | 0 | 0 | 0 | 0 | 3 | 0 | 0 | 0 | 2 | 0 |
| M16 | 203 | 29 | 0 | 0 | 0 | 8 | 3 | 5 | 2 | 1 | 0 | 4 | 0 | 2 | 0 | 0 | 1 | 0 | 1 | 0 | 0 | 0 | 0 | 0 |
| M1 | 491 | 22 | 0 | 0 | 1 | 4 | 0 | 3 | 3 | 0 | 0 | 5 | 0 | 1 | 0 | 0 | 0 | 0 | 0 | 0 | 0 | 0 | 3 | 0 |
| no cluster | 9336 | 265 | 8 | 6 | 7 | 85 | 13 | 31 | 16 | 8 | 14 | 7 | 0 | 14 | 0 | 0 | 2 | 3 | 5 | 4 | 1 | 3 | 15 | 7 |

#### Supplementary Table 4. KZFP members of the intronic TE module

KZFP members belonging to the intronic TE module N1, and module N10 that is correlated with module N1.

| KZFP members in the intronic TE module N1 | chromosome | KZFP members in correlated module N10 | chromosome |
|---|---|---|---|
| PRDM7\|11105 | 16 | ZNF44\|51710 | 19 |
| ZNF169\|169841 | 9 | ZNF738\|148203 | 19 |
| ZNF226\|7769 | 19 | ZNF439\|90594 | 19 |
| ZNF266\|10781 | 19 | ZNF337\|26152 | 20 |
| ZNF26\|7574 | 12 | ZNF334\|55713 | 20 |
| ZNF354B\|117608 | 5 | ZNF662\|389114 | 3 |
| ZNF700\|90592 | 19 | ZNF83\|55769 | 19 |
| ZNF7\|7553 | 8 | ZNF493\|284443 | 19 |
| ZNF789\|285989 | 7 | ZNF211\|10520 | 19 |
| ZNF814\|730051 | 19 | ZNF682\|91120 | 19 |
| ZNF841\|284371 | 19 | | |

#### Supplementary Table 5. Gene correlated with L1HS 5' transcript level

The list of genes correlated with L1HS 5' transcript level in more than one tissue. Gene names, the tissue where the significant correlation was found, coefficient of the gene estimated from the best linear model, p-value for the gene coefficient, q-value, and partial eta-squared for the gene are reported.

#### Supplementary Table 6. Housekeeping genes

List of housekeeping genes plotted in Figure 5 as reference.

| Caracausi et al. 2017 [71] | Eisenberg et al. 2013 [72] |
|---|---|
| ACTG1 | C1orf43 |
| RPS18 | CHMP2A |
| POM121C | C15orf24 |
| MRPL18 | EMC7 |

| | |
|---|---|
| TOMM5<br>YTHDF1<br>TPT1<br>RPS27 | GPI<br>PSMB2<br>PSMB4<br>RAB7A<br>REEP5<br>SNRPD3<br>VCP<br>VPS29 |

#### Supplementary Figure 1. Effect of normalization on TE transcripts

Total TE derived transcript count before and after normalization. Left panel shows the total reads counted by TEtranscripts plotted against the library size (total reads in the fastq file). Right panel shows the normalized read counts after the normalization process described in the Methods, again plotted against the library size.

#### Supplementary Figure 2. Example cases of correction for intron retention.

Cases where intron retention leads to TE derived transcripts. From top to bottom, AluSx1_dup59209(chrY:21153222-21153521), L2a_dup21781(chr2:113980079-113981081), L1MA7_dup4297 (chr8:134015602-134015763). AluSx1 is embedded within an exon of gene *TTY14*, L2a is embedded in an intron of *PAX8*, L1MA7 is embedded in an intron of gene *TG*. In all three cases, read counts for the focal TEs were reduced to zero, and the reads mapping to these TEs did not contribute to the overall TE family count.

#### Supplementary Figure 3. Comparison of TE family expression between multi-mapped reads and uniquely mapped reads.

Family level TE transcript quantification based on uniquely mapped reads (Bowtie1) and multi-mapped read(STAR) for six different TE families. AluSx1, AluYa5, HERVK3-int, HERVK9-int, L1HS and LTR5_Hs.

#### Supplementary Figure 4 co-expression modules in the weighted gene co-expression network analysis

Co-expression modules identified through the weighted gene co-expression network analysis (WGCNA) is visualized for the breast tissue data. The four TE modules M8, M21, M38, M45 are marked with a red *. a) Relationship between the 61 modules. b) correlation between the 61 modules. c) topological adjacency matrix used to identify the 61 modules.

#### Supplementary Figure 5. Transcription factor binding on KZFP genes that are members of the intronic TE module N1.

a) Transcription Factor binding enriched upstream of the genes that members of the intronic TE module N1. b) transcription factors enriched and the cell line that was assayed. Transcription factor binding enrichment is calculated by the EnrichR platform, based on the ChIP-Seq data collected in the ENCODE and ChEA databases.

#### Supplementary Figure 6. Transcription factor binding on genes correlated with L1HS 5’

Transcription Factor binding enriched upstream of the genes that are positively correlated with L1HS 5’ transcript level. Transcription factor binding enrichment is calculated by the EnrichR platform, based on the ChIP-Seq data collected in the ENCODE and ChEA databases.

#### Supplementary Figure 7 past radiation therapy and intronic TE expression

a. Intronic TE module expression profile. and b. total normalized read counts of all gene and TE members in intronic TE module N1. Plotted against the past radiation therapy in thyroid tissue.

#### Supplementary Figure 8 ENCODE candidate regulatory element marks overlapped with TE expression and ZFP binding

a-b. Overlap of TE expression and KZFP binding with ENCODE Candidate Regulatory Element marks. c-d. Proportion of each category of TEs that are marked with ENCODE Candidate Regulatory Element marks. a. and c. are TE counts in gene regions (including introns and +- 1Kb of start and end of genes). b. and d. are TE counts in intergenic regions (+-1Kb away from start and end of genes).

Supplementary Figure 1

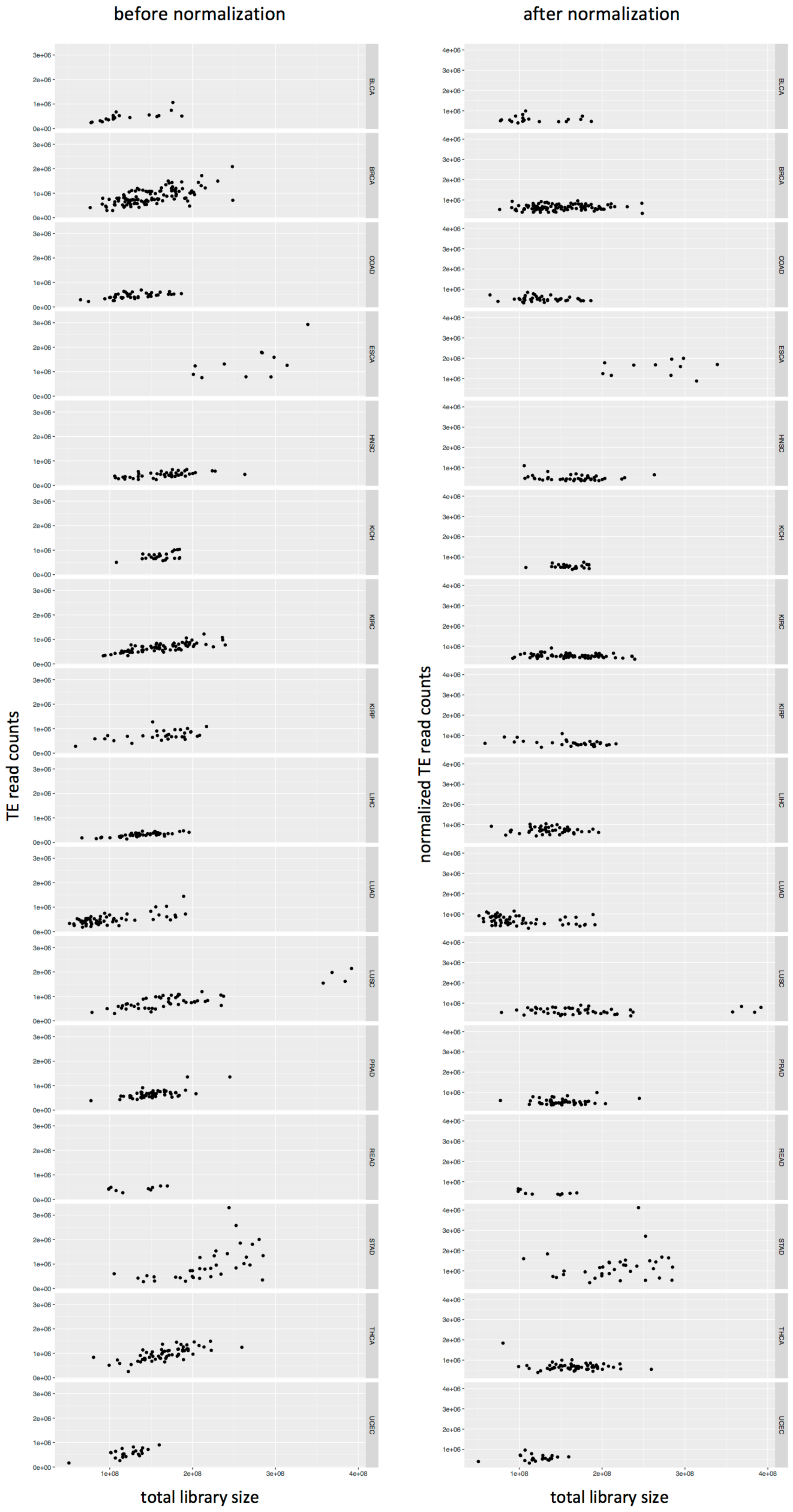

Supplementary Figure 2

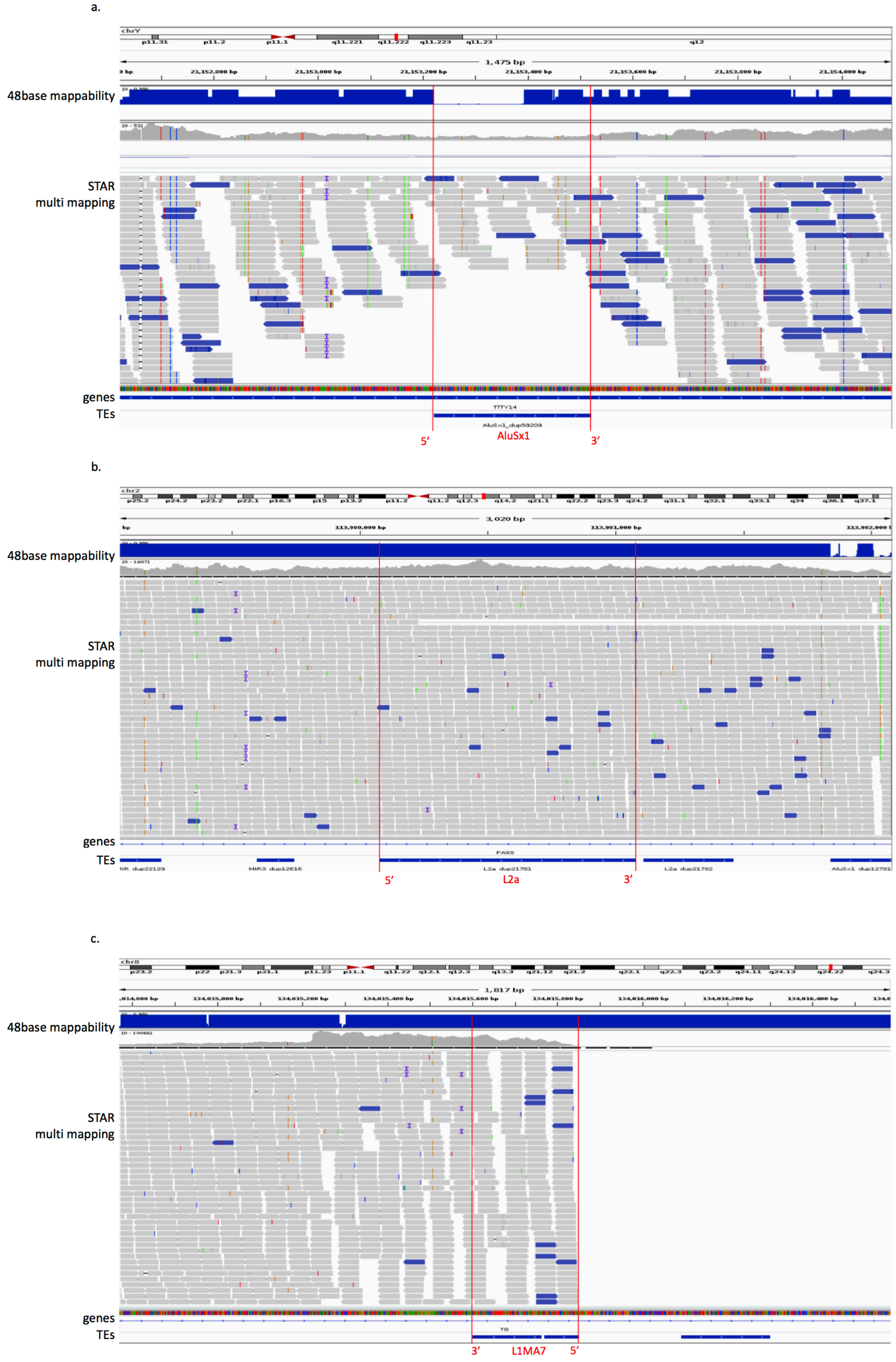

Supplementary Figure 3

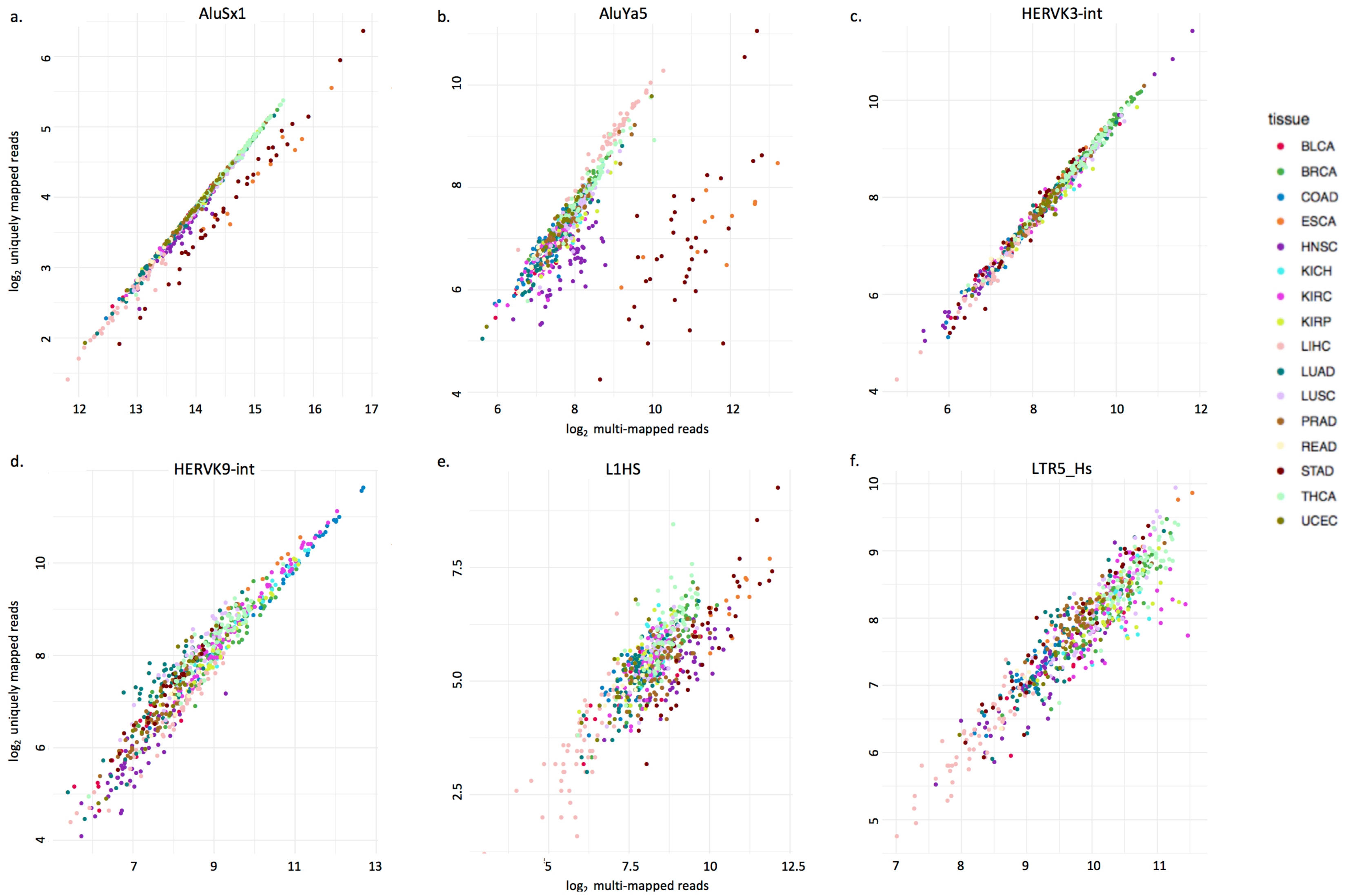

Supplementary Figure 4

a.

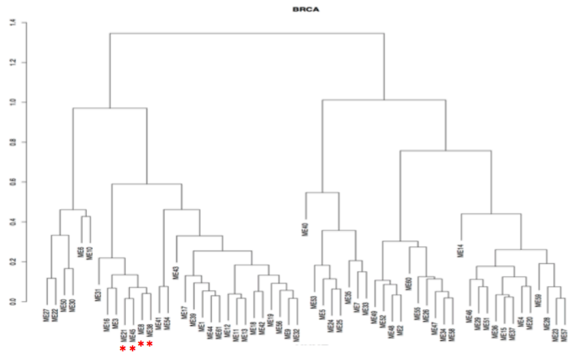

b.

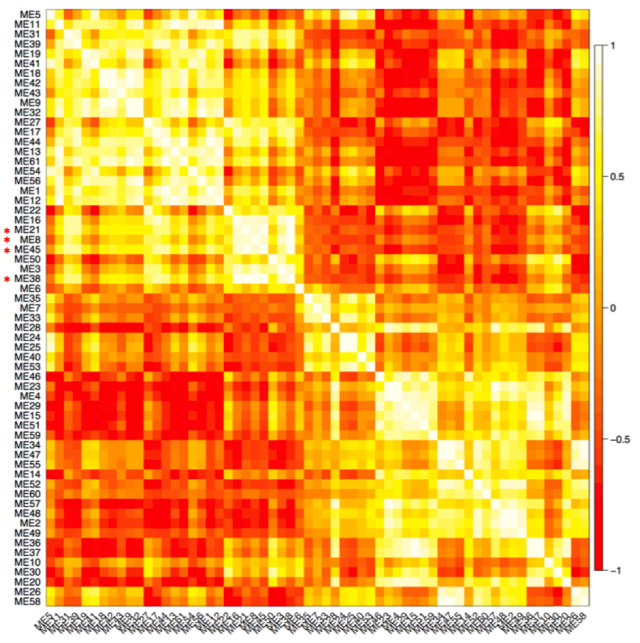

c.

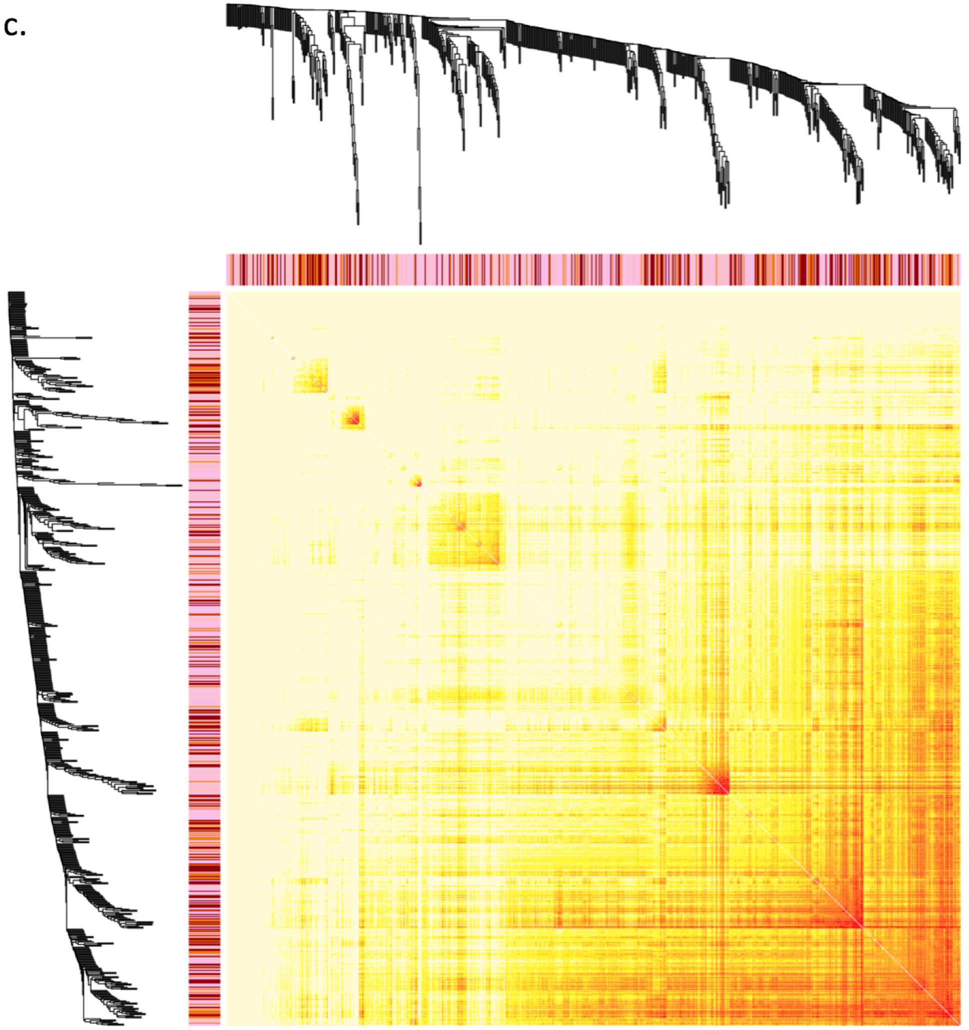

Supplementary Figure 5

a.

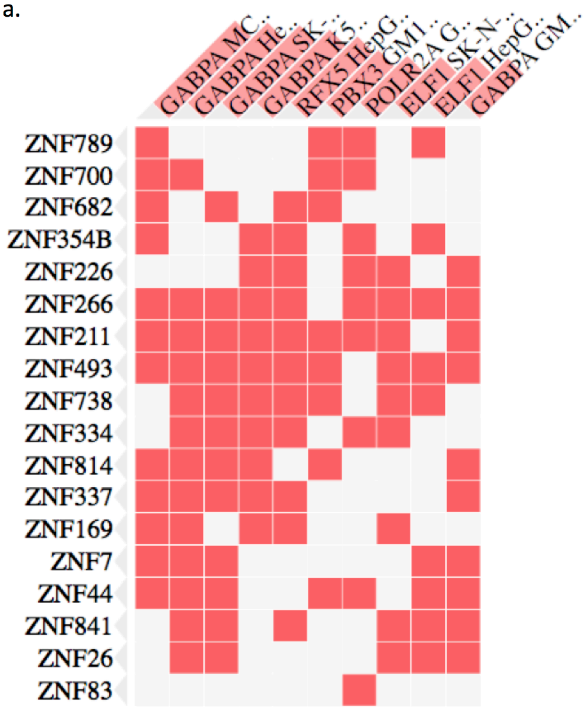

b.

GABPA_MCF-7
GABPA_HepG2
GABPA_SK-N-SH
GABPA_K562
RFX5_HepG2
PBX3_GM12878
POLR2A_GM18505
ELF1_SK-N-SH
ELF1_HepG2
GABPA_GM12878

Supplementary Figure 6

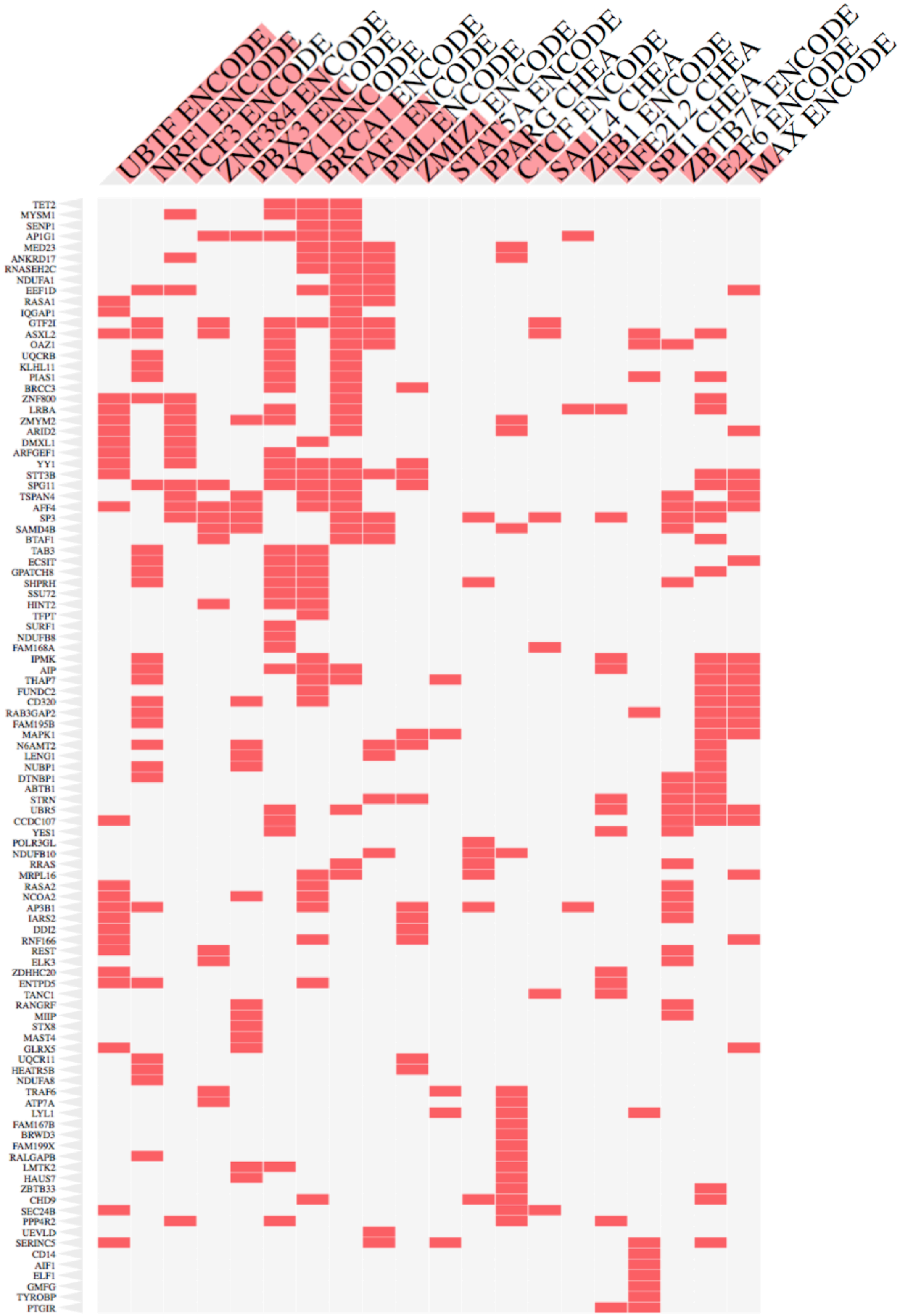

Supplementary Figure 7

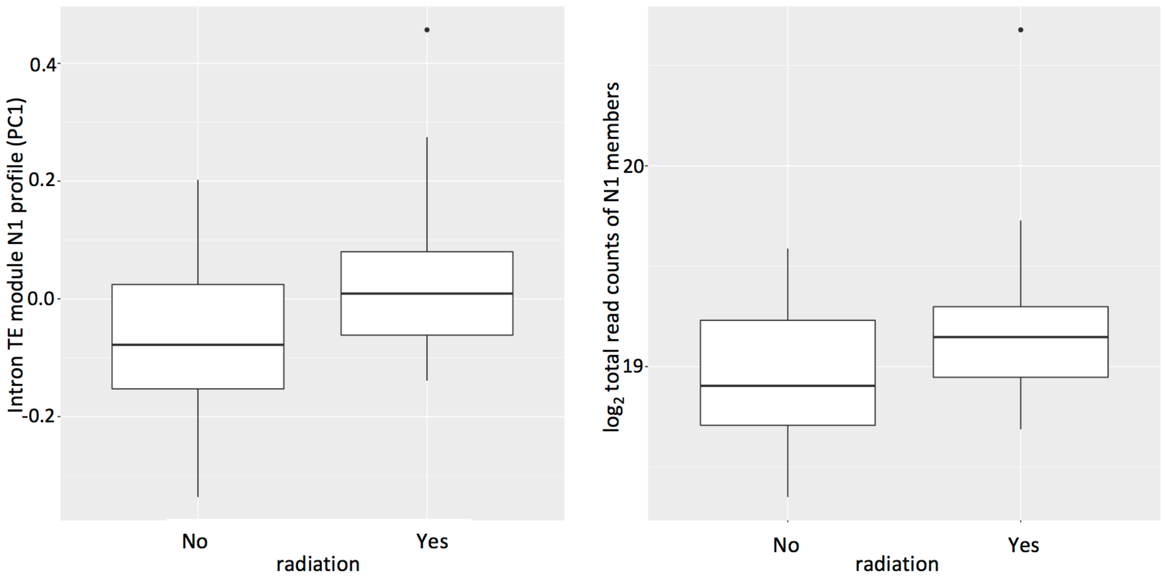

Supplementary Figure 8

a.

| ENCODE | RNA-seq | ZFP ChIP | overlap cnt | total cnt |
|---|---|---|---|---|
| | | ZNFbound | 11736 | 65358 |
| | expressed | ZNFunboud | 24364 | 352701 |
| | | ZNFbound | 18875 | 176105 |
| DNase+H3K4me3 | nonexpressed | ZNFunboud | 54469 | 1527856 |
| | | ZNFbound | 17735 | 65358 |
| | expressed | ZNFunboud | 89457 | 352701 |
| | | ZNFbound | 40271 | 176105 |
| DNase+H3K27ac | nonexpressed | ZNFunboud | 290920 | 1527856 |
| | | ZNFbound | 875 | 65358 |
| | expressed | ZNFunboud | 3230 | 352701 |
| | | ZNFbound | 2265 | 176105 |
| DNase+CTCFonly | nonexpressed | ZNFunboud | 14690 | 1527856 |

b.

| ENCODE | RNA-seq | ZFP ChIP | overlap cnt | total cnt |
|---|---|---|---|---|
| | | ZNFbound | 4634 | 51796 |
| | expressed | ZNFunboud | 9141 | 233016 |
| | | ZNFbound | 9157 | 186537 |
| DNase+H3K4me3 | nonexpressed | ZNFunboud | 37350 | 1902659 |
| | | ZNFbound | 13470 | 51796 |
| | expressed | ZNFunboud | 49592 | 233016 |
| | | ZNFbound | 39682 | 186537 |
| DNase+H3K27ac | nonexpressed | ZNFunboud | 282005 | 1902659 |
| | | ZNFbound | 1262 | 51796 |
| | expressed | ZNFunboud | 3694 | 233016 |
| | | ZNFbound | 3914 | 186537 |
| DNase+CTCFonly | nonexpressed | ZNFunboud | 24564 | 1902659 |

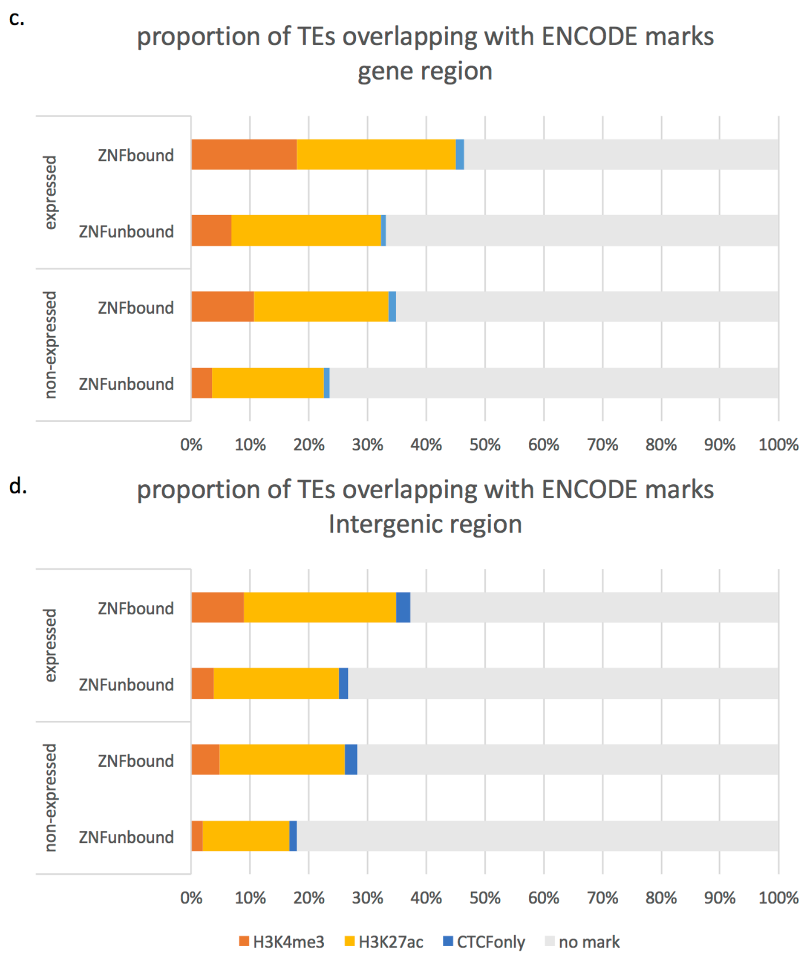
